# Persistently increased expression of PKMzeta and unbiased gene expression profiles identify hippocampal molecular traces of a long-term active place avoidance memory and ‘shadow’ proteins

**DOI:** 10.1101/2025.04.28.651143

**Authors:** Jiyeon Han, Alejandro Grau-Perales, Rayna M. Harris, Hsin-Yi Kao, Asit Pal, Juan Marcos Alarcon, Todd C. Sacktor, Stefano Martiniani, Hans A. Hofmann, Andre A. Fenton

## Abstract

Long-term memory formation transiently activates Ca^2+^-calmodulin kinase IIα and atypical protein kinase C isoform iota/lambda, whereas persistent activation of the other atypical PKC, protein kinase M zeta (PKMζ), together with its interacting partner, the scaffolding-protein KIBRA (*Wwc1*), are necessary for maintaining potentiated synapses and memory. Here, we use immediate-early gene (IEG) *Arc* activation during active place avoidance memory expression to tag memory-activated neurons with EYFP-ChR2. PKMζ immunohistochemistry identified which hippocampal synaptic pathways are persistently altered. EYFP-PKMζ colocalization persistently increases in the hippocampal trisynaptic pathway (dentate gyrus [DG]→CA3→CA1) tracing a one-month-old PKMζ engram. DG, CA3, and CA1 transcriptional profiling identifies that memory persistence correlates with upregulated immediate-early-genes *Arc*, *Fos*, and *NPas4* in DG, but not with *PKMζ* or most genes known to be crucial for LTP and memory. This rules out strong memory-related transcriptional regulation, but not regulation of mRNA translation or altered stability of “shadow proteins” like PKMζ that, despite being crucial for memory maintenance, evade detection by unbiased transcriptome profiling. In contrast, our method Correlation Signal Co-cluster Reduction (C-SCoRe) incorporates weak linear and non-linear gene correlations and highlights network interaction changes predicting memory, and related IEG and *Prkcz*/*Wwc1* expression. Manifold transcriptional relationships can reveal shadow molecular components of long-term memory.

## 1. Introduction

Learning about the significance of places empowers individuals to form and maintain adaptive and context-appropriate memories about locations associated with specific experiences, often for a lifetime. The cognitive and neural processes underlying place preference and avoidance have been examined in great detail (1), with much of this research focused on the rodent hippocampus, which is a thoroughly studied model system for investigating how spatial memory is formed and stored (2-4).

The active place avoidance task is used to assess spatial navigation and long-term memory in rodents. The task crucially depends on intact hippocampal function and causes month-long changes in hippocampal synaptic function at the level of the CA3-to-CA1 population of synapses and the medial entorhinal cortex-to-suprapyramidal dentate gyrus (DG) population of synapses (5-10). The task requires a rodent to explore a circular rotating arena while avoiding the location of a mild footshock. Place avoidance learning causes the transient activation of several long-term potentiation (LTP)-associated molecules in the dorsal hippocampus, including a number of kinases such as the persistent kinases Ca^2+^-calmodulin kinase IIα (CaMKII) and the atypical protein kinase C isoform PKC𝜄/λ for the acquisition of the conditioned avoidance (11-13). Importantly, these molecules are not critical for maintaining memory. In contrast, the other atypical PKC, protein kinase M zeta (PKMζ), an autonomously active kinase whose activity is regulated by its protein level (14), persistently increases and is necessary for long-term memory for at least a month (11, 15, 16), as shown across diverse brain regions, species, and memory paradigms (3, 17-31). Several input-specific populations of synapses in the hippocampus undergo synaptic changes, possibly to store memory and realize experience-dependent modifications of information processing (5, 8-10, 20, 32). However, which synaptic populations persistently change with experience remains an open question. One hypothesis is that these changes primarily occur in those synapses that are relevant for the maintenance of the memory, in which case we can predict that they will be expressed in distinct synaptic populations within the different hippocampal subfields (10, 32, 33). An alternative hypothesis posits that changes occur everywhere in a non-specifically structured organization (34, 35). The substantial recurrent connectivity of the CA3 subfield has motivated the theory that it might act as a storage site with robust memory-related synaptic plasticity (36-38). Studies of hippocampal neurons that express location-specific place cell discharge identified the importance of the CA3-to-CA1 synapses in memory storage and recall. In contrast, the entorhinal cortex layer III-to CA1 temporoammonic (TA) synaptic terminals are thought to be relevant for encoding momentary information and computations (36, 39-41). Inputs from layer II of the lateral entorhinal cortex are thought to provide contextual information, while those from layer II of the medial entorhinal cortex provide spatial signals. These pathways differentially target dendritic compartments in the supra- and infra-pyramidal blades of dentate gyrus (42-44). The specific synaptic organization of the hippocampal circuit is likely to constrain, and possibly predict, the functional organization of hippocampal information processing.

Molecules such as PKMζ that are crucial for memory maintenance, interact with other effector and regulatory molecules in multiple ways that have yet to be completely identified, but likely are best characterized as a network of interactions abstracted from the list of the molecular components (11, 13, 28, 45-56). Although the complete network of memory-mediating molecular interactions that maintain persistent place avoidance memory is unknown, biochemical studies have shown that PKMζ associates with the kidney and brain protein (KIBRA/WWC1) and potentiates synaptic transmission through the action of N-ethylmaleimide-sensitive factor (NSF), which in turn increases the insertion of GluA2 subunit-containing α-amino-3-hydroxy-5-methyl-4-isoxazolepropionic acid (AMPA)-type glutamate ionotropic receptors into postsynaptic sites (45, 55). This process coincides and integrates with a reduction in NUMB-mediated AMPA receptor endocytosis coordinated by protein interacting with C-kinase 1 (PICK1) from the postsynaptic sites that are maintained by CaMKII-activity/structure (45, 56-58). It is not known, however, whether these processes are largely regulated at the post-transcriptional level, including regulation of mRNA translation and protein stability, or whether they are accompanied by changes in the mRNA levels of these genes, possibly in opposing directions (e.g., PICK1 and NUMB functions are anti-related to the memory-promoting cell biological functions of PKMζ, KIBRA, NSF, CaMKII, and GluA2). Furthermore, LTP-induction is associated with transient upregulation of PKC𝜄/λ, the other atypical PKC, whereas genetic deletion of the PKMζ gene *Prkcz*, results in LTP- and memory-triggered prolongation of compensatory upregulation of PKC𝜄/λ and the conventional isoform PKC*β*1 (11). It is also unclear how this PKMζ-dependent memory-maintenance system interacts with other molecular pathways that have been hypothesized to play a role in the early phases of memory formation. These include mTOR, a master regulator of translation, insulin-like growth factor 2 (IGF2), which promotes the strengthening of memory, likely by an independent process (59, 60), or Fragile X Messenger Ribonucleoprotein 1 (FMR1), the loss of which can lead to dysregulated mRNA translation, altered synaptic function, abnormal protein synthesis-dependent synaptic plasticity, and poorly coordinated neural discharge (61-63). These findings suggest that changes in both mRNA expression as well as post-transcriptional regulation of genes involved in synaptic plasticity underlie memory maintenance. Modern genomic techniques such as transcriptome profiling have made it possible to examine the molecular underpinnings of plasticity in animal behavior and can be profitably used to test specific hypotheses about the molecular underpinnings of memory maintenance as well as discover novel pathways for subsequent experimental analysis (64).

In the present study, we used active place avoidance as an exemplar of long-term memory maintenance in two separate experiments. Experiment 1 used immunohistochemistry to test the hypothesis that PKMζ exerts its memory-maintaining role by localizing to specific cellular sub-compartments of EYFP-tagged memory cells in the hippocampal subfields. Experiment 2 used RNA-seq in specific hippocampal subfields to test the hypothesis that the transcriptional regulation of the PKMζ system, along with several other candidate genes, are critical for memory maintenance in the active place avoidance paradigm. Lastly, we mined the RNA-seq data obtained from DG, CA3, and CA1 in search of covariant molecular networks predictive of memory and PKMζ gene expression levels, used here as an exemplar of proteins with crucial functions, such as in memory maintenance, but which are differentially expressed at very low levels, limiting their detection by standard differential expressed gene (DEG) or linear correlation analysis. We refer to this class of molecules as the ‘shadow proteome,’ which is distinct from the so-called ‘dark proteome’ of proteins originating from unknown open-reading frames and DNA sequences (65). Using methods to measure both linear and non-linear correlations, we developed a new analysis pipeline Correlation Signal Co-Cluster Reduction (C-SCoRe) to identify how molecular interactions change with a perturbation, in this case memory training. C-SCoRe exploits the within-subject correlations amongst gene expression values instead of their abundance. As used, it selects genes related to behavioral measures of memory, and projects the large network of gene-gene interaction changes into low-dimensional spaces defined by their covariance. C-SCoRe was able to predict the observed relationship between PKMζ gene expression and memory persistence, validating its utility and demonstrating the feasibility of detecting shadow proteins and potentially identifying novel molecular interactions and pathways associated with memory maintenance, even when levels of differential gene expression cannot be detected.

## 2. Results

### 2.1. Experiment 1 – Tracing Memory

#### 2.1.1. PKMζ protein is persistently increased for at least 30 days in memory-tagged cells

The Trained and Untrained ArcCreER^T2^ x ChR2-EYFP TRAP2 transgenic mice (Figure 1A) do not differ in entering the shock zone soon after being placed on the arena during pretraining (Figure 1B,C; t_7_ = 0.42, p = 0.7), but only the Trained mice learn to delay entering the shock zone for several minutes by the end of training (t_7_ = 4.18, p = 0.006). This conditioned avoidance persists and is expressed a month later in the retention test with no shock (Trained: 90 ± 51 s, Untrained: 5 ± 4 s). Despite the absence of shock, the time to enter the location of shock for the second time is 254 ± 141 s in the Trained group and only 26 ± 17 s in the Untrained group (t_7_ = 3.60, p = 0.01), confirming place avoidance memory is maintained for a month in the Trained mice (Figure 1C).

**Figure 1.**
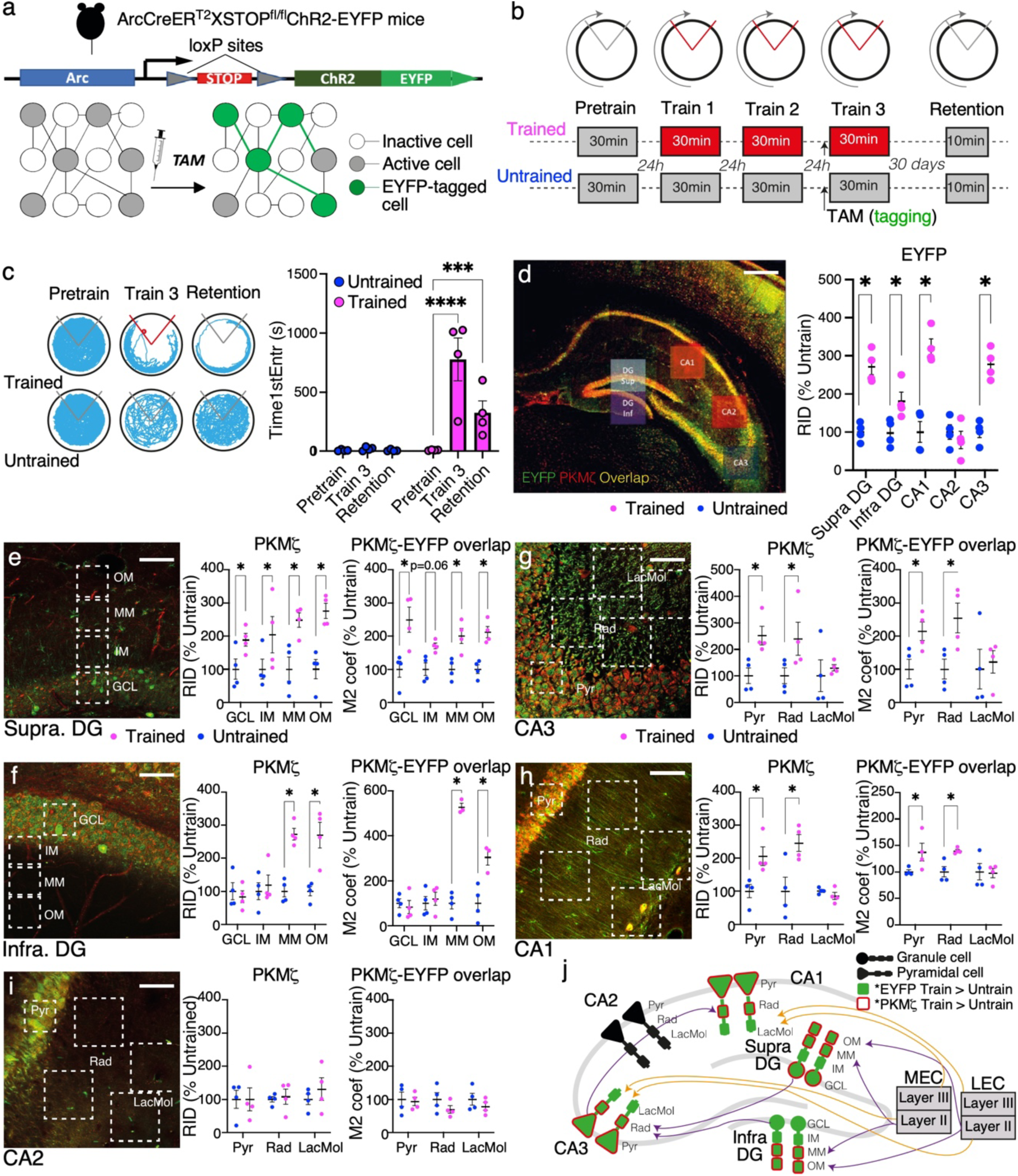
Persistently elevated PKMζ protein expression in dorsal hippocampus subcellular compartments that trace the trisynaptic pathway circuit after 1-mo active place avoidance memory. a) Schematic of the TRAP2 mouse that was engineered to permanently express the EYFP-ChR2 tag in neurons that activate the immediate early gene *Arc* that is expressed primarily in excitatory neurons. b) Schematic of the behavioral experimental design. After learning an active place avoidance, EYFP-ChR2 tagging was initiated by systemic injection of 4OH-tamoxifen (TAM) 30 min prior to the third training session of the place avoidance memory and untrained control groups. Thirty days later, memory retention is tested prior to euthanasia and collection of tissue for PKMζ immunohistochemistry (n’s = 4). c) Example of spatial exploration on the arena for exemplar Trained and Untrained mice to illustrates the conditioned behavior, the summary plot quantitates the active place avoidance. d) Merged EYFP-ChR2 (green) and PKMζ (red) immunofluorescent image of dorsal hippocampus of a Trained mouse, with ROIs for quantifying Training 3 memory activation by expression of EYFP-ChR2 and memory retention by PKMζ expression computed as the raw integrated density (RID), each normalized by the control average. e) Summary of quantitation of EYFP expression. f-i) Example ROI samples targeting specific subcellular tissue compartments of the dorsal hippocamous circuit. Plots summarize control-normalized RID quantitation of PKMζ expression in each subcompartment and co-expression of EYFP and PKMζ, estimated by the control-normalized Mander’s overlap coefficient (M2), which computes the ratio of PKMζ expression (RID) at image elements that express EYFP. j) Graphical summary illustrating the hippocampal circuit compartments that are persistently altered by increased PKMζ protein expression 1 month after memory training. Scale bars in panel d: 300 µm; e-I: 50 µm.

The EYFP label is detected throughout the principal cells of Ammon’s horn, including the *strata radiatum* and *lacunosum moleculare* dendritic compartments and the granule cell and the molecular layers of the supra-DG and infra-DG of both Trained and Untrained mice (Figure 1D). We first correlate the somatic EYFP signal (in *str. pyramidale* for CA1, CA2, and CA3 and in the granule cell layer for DG) with the dendritic compartment signals to assess whether the EYFP label is distributed across the *strata* that define synaptic compartments. Within each hippocampal subfield, we observe strong positive correlations between somatic EYFP expression and each of the corresponding dendritic compartments (0.34 ≥ r’s ≤ 0.90; 0.89 < t_6_’s < 5.1, 0.002 ≥ p’s ≤ 0.4). These correlations are all significant (p’s < 0.05) except between the granule cell layer and the adjacent inner molecular layer of infra-DG (r = 0.54, t_6_ = 1.6, p = 0.2) and between the pyramidal cell layer of CA3 and the distant *str. lacunosum moleculare* compartment (r = 0.34, t_6_ = 0.89, p = 0.4).

We next determine whether memory training causes elevated EYFP expression in each hippocampal subregion. Relative to control expression values (Figure 1D), EYFP expression is significantly increased in the Trained group at CA1 (t_6_ = 2.81, p = 0.03), CA3 (t_6_ = 3.80, p = 0.008), supra-DG (t_6_ = 8.61, p <0.001), and infra-DG (t_6_=2.58, p = 0.04), but not at CA2 (t_6_ = 1.20, p = 0.3).

We can now investigate which EYFP-tagged synaptic populations persistently increase PKMζ expression after memory training as an indicator of persistently potentiated memory-related synapses. Compared to Untrained, PKMζ is significantly increased throughout the Trained supra-DG (Figure 1E). Although Mander’s M1 coefficients do not significantly differ (all p’s > 0.1), the M2 coefficient estimating PKMζ-EYFP overlap exclusively in EYFP expressing regions is significantly increased in Trained mice at the granule cell layer (t_6_ = 2.65, p = 0.02), the middle (t_6_ = 3.11, p = 0.01), and the outer molecular layers (t_6_ = 4.11, p = 0.003), but not at the inner molecular layer (t_6_ = 2.24, p = 0.06), which only reaches 1-tailed significance. This suggests that synapses of the outer two-thirds of the molecular layer of supra-DG are persistently potentiated a month after memory training, coinciding with where the inputs from LECII and MECII terminate in supra-DG, consistent with electrophysiological observations in freely-behaving mice (5).

The corresponding analysis of infra-DG finds Trained versus Untrained enhancement of PKMζ only at the middle and outer molecular layers (Figure 1F) and although there are also no group differences in M1 coefficients, the M2 measure of PKMζ-EYFP overlap is significantly increased in Trained compared to Untrained mice at the middle (t_6_ =14.4, p < 0.01) and outer molecular layers (t_6_ = 4.15, p < 0.01), but not elsewhere (Figure 1F).

CA3 labeling for PKMζ is significantly increased in Trained compared to Untrained *str. pyramidale* (t_6_ = 3.28, p = 0.01) and the *str. radiatum* (t_6_ = 3.18, p = 0.01), but not at *str. lacunosum moleculare* (t_6_ = 0.49, p = 0.6). While we do not find any differences between groups in the M1 coefficient (all p’s > 0.4), this coincides with increase of the M2 overlap coefficient at *str. pyramidale* (t_6_ = 2.80, p = 0.03), and *str. radiatum* (t_6_ = 2.59, p = 0.04), but not *str. lacunosum moleculare* (t_6_ = 0.32 p = 0.8), suggesting persistent potentiation of synaptic inputs from DG, but not entorhinal cortex in memory-tagged CA3 (Figure 1G).

PKMζ in the CA1 subfield is significantly increased in Trained compared to Untrained mice at *str. pyramidale* (t_6_ = 3.00, p = 0.01), and *str. radiatum* (t_6_ = 3.63, p = 0.01), but not *str. lacunosum moleculare* (t_6_ = 1.24, p = 0.3; Figure 1H). Similar to CA3, we do not observe group differences in the M1 coefficient (p’s > 0.1), but the M2 coefficient is increased at *str. pyramidale* (t_6_ = 2.80, p = 0.03), and *str. radiatum* (t_6_ = 2.59, p = 0.04), but not *str. lacunosum moleculare* (t_6_ = 0.32, p = 0.8) of the memory-tagged cells, which replicates a prior report (66) and is consistent with long-term memory training causing persistent electrophysiological potentiation of the Schaffer collateral-commissural population of synapses (10).

Within CA2, we do not find any differences in the expression of PKMζ in any of the synaptic compartments (Figure 1I; p’s > 0.4), the M1 (p’s ≥ 0.2), and the M2 coefficients (p’s ≥ 0.3) in any of the synaptic compartments of the CA2 region, consistent with observing no evidence of training-associated *Arc* activation.

The overall hippocampal pattern of changes in PKMζ expression in memory-tagged cells is remarkable. One month after memory training, the PKMζ increase traces the synaptic compartments of the trisynaptic pathway through the hippocampal circuit (Figure 1J).

#### 2.1.2. Results motivating Experiment 2

These findings are consistent with the hypothesis that place avoidance training causes synaptic potentiation in the hippocampal trisynaptic “model” pathway thought to be associated with memory, but not in the direct “data” inputs from entorhinal cortex thought to be associated with sensory information processing (39, 40, 67, 68). This PKMζ memory trace is selectively observed in the neurons that activate *Arc* during expression of the established place avoidance memory, not merely during learning (Figure 1), in line with observations that memory training increases PKMζ in CA1 (15, 66) and changes synaptic function selectively at CA1 *str. radiatum* (10) as well as Supra-DG, which can be measured as early as after 30 min of memory training (5), see also (32). The observed group differences in PKMζ predict that mice with better memory will express more PKMζ in the memory-tagged cells. Despite there being only four Trained mice, large Pearson correlations are observed between the M2 coefficients of PKMζ in EYFP-expressing cells and the time to first enter into the shock zone on Trial 3 when memory was tagged. Correcting for the two calculations (M1, and M2), the M2 coefficient is significant at the middle molecular layer where the medial perforant path terminates in the supra-DG (r = 0.98, t_2_ = 7.0, p = 0.02), the site where Chung et al. recorded memory-induced synaptic plasticity in freely-behaving mice for up to 60 days; but the correlation is not significant at infra-DG (r = 0.93, t_2_ = 3.7, p = 0.06), mirroring the electrophysiological observations (5, 69). There is strong evidence that PKMζ protein expression is persistently and detectably elevated at synaptic populations, specifically in the ChR2-EYFP-tagged neurons that are an enriched sample of memory-related cells. Lastly, prior work showed that the training-induced persistent increase of PKMζ in memory-tagged cells and the synaptic potentiation at CA1 *str. radiatum*, are both indistinguishable whether or not memory retention is tested (10, 15, 66).

### 2.2. Experiment 2 – Transcriptional Profiling of Memory

#### 2.2.1. Characterizing memory-related behavior

We first analyze active place avoidance behavior in the memory-trained and yoked-control groups of wild-type mice, as a prerequisite for investigating transcriptional profiles related to memory (Figure 2). During pretraining, when the shock was off, there are no differences between the groups, as expected (Table 1). Avoidance is evaluated in multiple ways, including as a reduction in the number of entrances into the shock zone (Figure 3A), an increase in the time to first enter the shock zone (Figure 3B), and a decreased likelihood of being in the shock zone (Figure 3C). As expected, the memory-trained animals show a robust conditioned avoidance whereas the yoked-controls do not, although corresponding pairs of Trained and Control mice experience identical physical conditions (Figure 2B). These observations are confirmed by two-way ANOVAs, with significant effects of training treatment, trial, and their interaction (Table 1).

**Figure 2.**
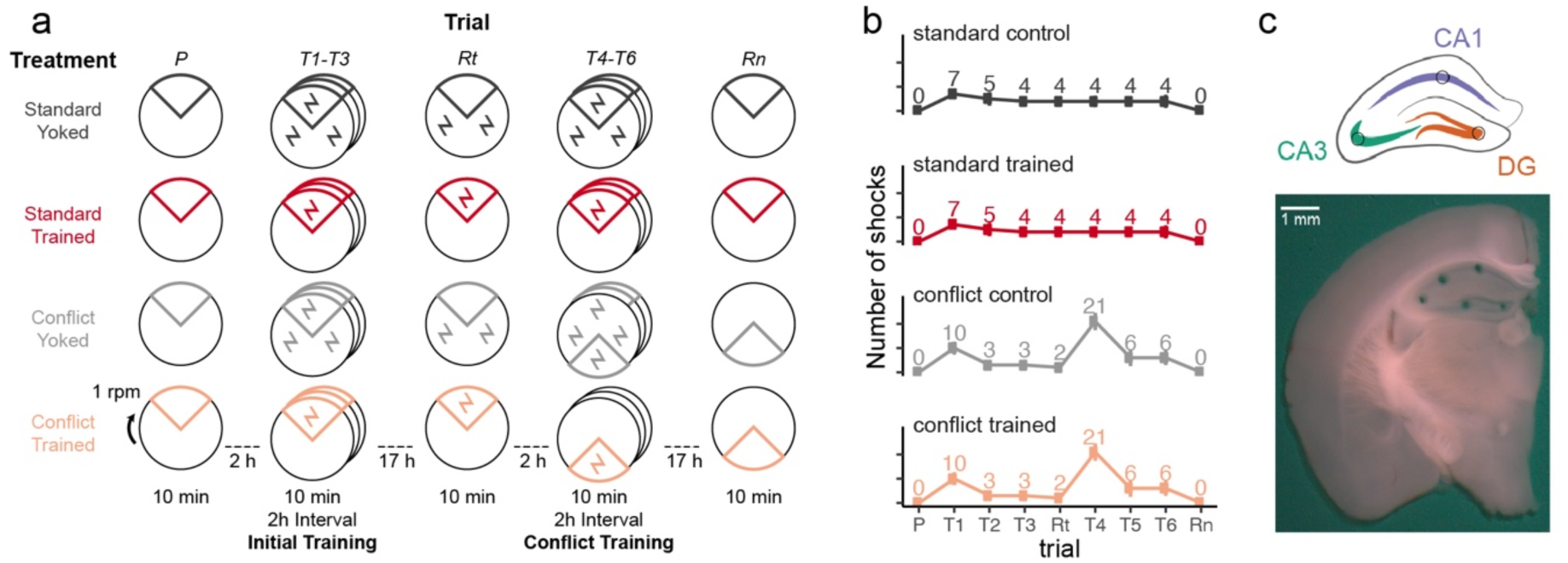
Experimental Design. a) WT mice are assigned to one of four groups: standard-trained (red, n=8), standard-yoked (dark grey, n=8), conflict-trained (peach, n=9), or conflict-yoked (light grey, n=9). Mice were placed on the rotating arena for 10-min trials separated by 2-h or overnight (∼17 h) intervals. Behavior was tracked during the pretraining (P), initial training (T1-T3), retest (Rt), conflict training (T4-T6), and retention (Rn) trials. In the trial schematics, the 60° sector represents the shock zone or for the yoked groups, the equivalent region that is used for evaluating place avoidance. Whereas trained mice receive shocks only in the shock zone, yoked mice experience shock throughout the environment. b) Standard-trained and standard-yoked mice received an identical number of shocks (and time series), with the most shocks occurring on trial 1. Conflict-trained and conflict-yoked mice received an identical number of shocks (and time series), with the most shocks occurring on trial 4, the first trial of conflict training. c) A schematic representation and a photo show the size and location of tissue samples collected from the dorsal hippocampus for RNA-sequencing. Tissues from five subfields (CA1, CA2, CA3, CA4, DG) were collected, but only the DG (orange), CA3 (teal), and CA1 (purple) subfields were processed for RNA-seq.

**Figure 3.**
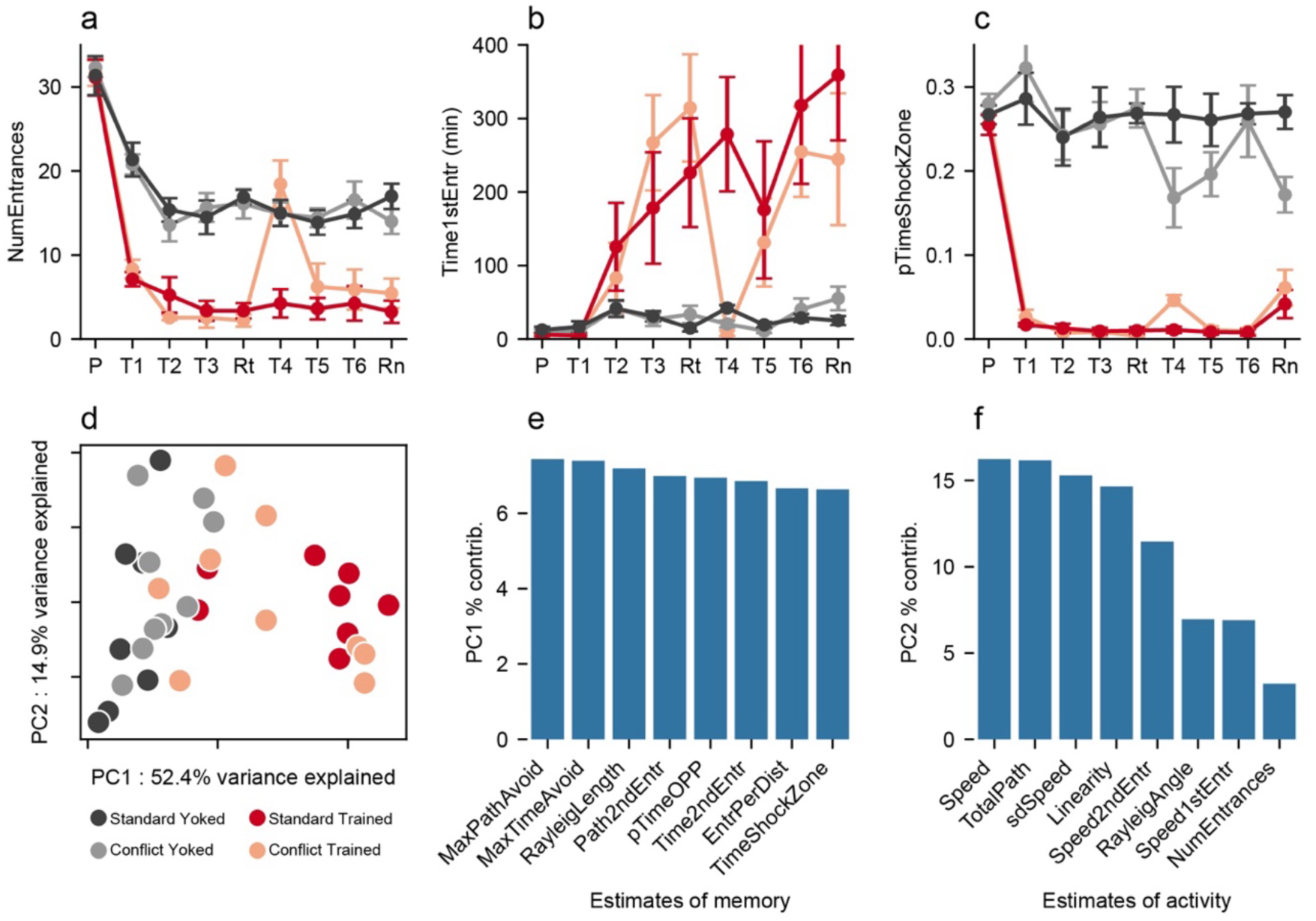
Characterizing and comparing behavior amongst the groups. Standard-trained and conflict-trained groups rapidly learned to avoid the shock zone, whereas the yoked control groups did not avoid the corresponding region measured by a) reduction in the number of entrances into the shock zone, b) increased latency to enter the shock zone for the first time on each trial, and c) reduction in the proportion of time spent in the shock zone and its equivalent. d) PCA performed on the 26 measures of behavior during retention, results in PC1 that distinguishes the Trained and Control groups just prior to sacrifice when physical conditions are identical for all the mice because there is no shock. Large dots indicate group averages on the retention test while transparency is highest for pretraining and lowest for retention. Parameter loadings for e) PC1 and f) PC2 indicate that PC1 is comprised of estimates of place avoidance and PC2 estimates of locomotor behavior. Comparison statistics are provided in Table 1. MaxPathAvoid = maximum distance walked without entering the shock zone; MaxTimeAvoid = maximum time (s) spent out of the shock zone; RayleighLength = the Rayleigh vector length estimating the angular bias of the animal’s dwell probability; Path2ndEntr = distance (m) walked before entering the shock zone twice; pTimeOPP = proportion of time spent in the quadrant opposite from the target quadrant; Time2ndEntr = amount of time it took the animal to enter the shock zone for the second time; EntrPerDist = number of entrances per meter walked; TimeShockZone = total time (s) spent in the shock zone; Speed = Average speed of the animal throughout the session; TotalPath = total distance (m) walked; sdSpeed = standard deviation of the speed estimates; Linearity = straightness of locomotion, estimated every 2 seconds as the ratio of the end-to-end Euclidean distance divided by the integral of the path segments; Speed2ndEntr = speed when entering the shock zone the second time; RayleighAngle = angle of the vector describing where the animal spent most of its time (the angle describes the particular preferred location. Ranges 0-360); Speed1stEntr = speed when entering the shock zone the first time; NumEntrances = total number of entrances into the shock zone.

**Table 1.**
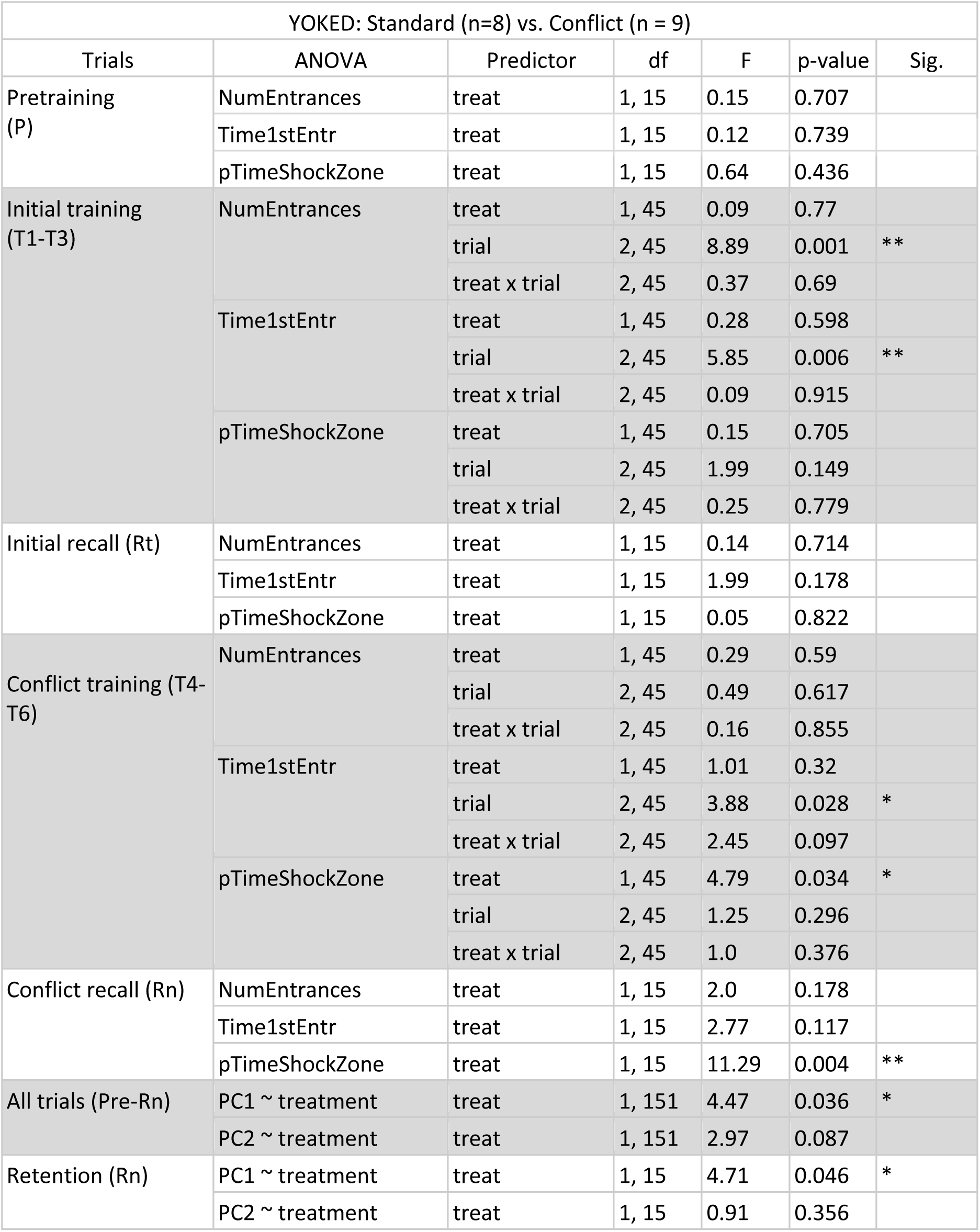
One-way and two-way ANOVA results of place avoidance behavior analysis of the yoked groups. One-way ANOVAs were used to determine the effects of training treatment on behavior during a single trial, including the pretraining and recall trails. Two-way ANOVAs were used to determine the effects of treatment, training, and interaction across three trials, including the initial training session and the conflict training session. Analyses were performed separately for the trained and yoked groups. Key: NumEntrances = total number of entrances into the shock zone; Time1stEntr = time (min) passes before entering the shock zone the first time; pTimeShockZone = proportion of time spent in the

| YOKED: Standard (n=8) vs. Conflict (n = 9) |  |  |  |  |  |  |
| --- | --- | --- | --- | --- | --- | --- |
| Trials | ANOVA | Predictor | df | F | p-value | Sig. |
| Pretraining (P) | NumEntrances | treat | 1, 15 | 0.15 | 0.707 |  |
|  | Time1stEntr | treat | 1, 15 | 0.12 | 0.739 |  |
|  | pTimeShockZone | treat | 1, 15 | 0.64 | 0.436 |  |
| Initial training (T1-T3) | NumEntrances | treat | 1, 45 | 0.09 | 0.77 |  |
|  |  | trial | 2, 45 | 8.89 | 0.001 | ** |
|  |  | treat x trial | 2, 45 | 0.37 | 0.69 |  |
|  | Time1stEntr | treat | 1, 45 | 0.28 | 0.598 |  |
|  |  | trial | 2, 45 | 5.85 | 0.006 | ** |
|  |  | treat x trial | 2, 45 | 0.09 | 0.915 |  |
|  | pTimeShockZone | treat | 1, 45 | 0.15 | 0.705 |  |
|  |  | trial | 2, 45 | 1.99 | 0.149 |  |
|  |  | treat x trial | 2, 45 | 0.25 | 0.779 |  |
| Initial recall (Rt) | NumEntrances | treat | 1, 15 | 0.14 | 0.714 |  |
|  | Time1stEntr | treat | 1, 15 | 1.99 | 0.178 |  |
|  | pTimeShockZone | treat | 1, 15 | 0.05 | 0.822 |  |
| Conflict training (T4-T6) | NumEntrances | treat | 1, 45 | 0.29 | 0.59 |  |
|  |  | trial | 2, 45 | 0.49 | 0.617 |  |
|  |  | treat x trial | 2, 45 | 0.16 | 0.855 |  |
|  | Time1stEntr | treat | 1, 45 | 1.01 | 0.32 |  |
|  |  | trial | 2, 45 | 3.88 | 0.028 | * |
|  |  | treat x trial | 2, 45 | 2.45 | 0.097 |  |
|  | pTimeShockZone | treat | 1, 45 | 4.79 | 0.034 | * |
|  |  | trial | 2, 45 | 1.25 | 0.296 |  |
|  |  | treat x trial | 2, 45 | 1.0 | 0.376 |  |
| Conflict recall (Rn) | NumEntrances | treat | 1, 15 | 2.0 | 0.178 |  |
|  | Time1stEntr | treat | 1, 15 | 2.77 | 0.117 |  |
|  | pTimeShockZone | treat | 1, 15 | 11.29 | 0.004 | ** |
| All trials (Pre-Rn) | PC1 ~ treatment | treat | 1, 151 | 4.47 | 0.036 | * |
|  | PC2 ~ treatment | treat | 1, 151 | 2.97 | 0.087 |  |
| Retention (Rn) | PC1 ~ treatment | treat | 1, 15 | 4.71 | 0.046 | * |
|  | PC2 ~ treatment | treat | 1, 15 | 0.91 | 0.356 |  |

**Table 2.** One-way and two-way ANOVA results of place avoidance behavior analysis of the trained groups.

| TRAINED: Standard (n=8) vs. Conflict (n = 9) |  |  |  |  |  |  |
| --- | --- | --- | --- | --- | --- | --- |
| Trials | ANOVA | Predictor | df | F | p-value | Sig. |
| Pretraining (P) | NumEntrances | treat | 1, 15 | 0.04 | 0.837 |  |
|  | Time1stEntr | treat | 1, 15 | 0.96 | 0.343 |  |
|  | pTimeShockZone | treat | 1, 15 | 0.51 | 0.485 |  |
| Initial training (T1-T3) | NumEntrances | treat | 1, 45 | 0.6 | 0.444 |  |
|  |  | trial | 2, 45 | 8.9 | 0.001 | ** |
|  |  | treat x trial | 2, 45 | 1.28 | 0.288 |  |
|  | Time1stEntr | treat | 1, 45 | 0.15 | 0.697 |  |
|  |  | trial | 2, 45 | 9.33 | 0.0 | *** |
|  |  | treat x trial | 2, 45 | 0.85 | 0.433 |  |
|  | pTimeShockZone | treat | 1, 45 | 0.06 | 0.805 |  |
|  |  | trial | 2, 45 | 4.01 | 0.025 | * |
|  |  | treat x trial | 2, 45 | 1.07 | 0.351 |  |
| Initial recall (Rt) | NumEntrances | treat | 1, 15 | 0.97 | 0.341 |  |
|  | Time1stEntr | treat | 1, 15 | 0.72 | 0.411 |  |
|  | pTimeShockZone | treat | 1, 15 | 2.38 | 0.144 |  |
| Conflict training (T4-T6) | NumEntrances | treat | 1, 45 | 10.7 | 0.002 | ** |
|  |  | trial | 2, 45 | 5.68 | 0.006 | ** |
|  |  | treat x trial | 2, 45 | 4.61 | 0.015 | * |
|  | Time1stEntr | treat | 1, 45 | 4.56 | 0.038 | * |
|  |  | trial | 2, 45 | 2.56 | 0.089 |  |
|  |  | treat x trial | 2, 45 | 1.51 | 0.233 |  |
|  | pTimeShockZone | treat | 1, 45 | 13.31 | 0.001 | ** |
|  |  | trial | 2, 45 | 14.54 | 0.0 | *** |
|  |  | treat x trial | 2, 45 | 9.62 | 0.0 | *** |
| Conflict recall (Rn) | NumEntrances | treat | 1, 15 | 0.96 | 0.342 |  |
|  | Time1stEntr | treat | 1, 15 | 0.82 | 0.381 |  |
|  | pTimeShockZone | treat | 1, 15 | 0.49 | 0.495 |  |
| All trials (Pre-Rn) | PC1 ~ treatment | treat | 1, 151 | 1.16 | 0.283 |  |
|  | PC2 ~ treatment | treat | 1, 151 | 0.0 | 0.980 |  |
| Retention (Rn) | PC1 ~ treatment | treat | 1, 15 | 1.48 | 0.243 |  |
|  | PC2 ~ treatment | treat | 1, 15 | 0.0 | 0.959 |  |

**Table 3.** One-way and two-way ANOVA results of place avoidance behavior analysis comparing merged yoked and trained groups. The predictor term ‘train’ indicates the two binary training conditions (Yoked vs. Trained).

| YOKED (n=17) vs. TRAINED (n=17) |  |  |  |  |  |  |
| --- | --- | --- | --- | --- | --- | --- |
| Trials | ANOVA | Predictor | df | F | p- | Sig. |
| Pretraining (P) | NumEntrances | train | 1, 32 | 0.07 | 0.789 |  |
|  | Time1stEntr | train | 1, 32 | 1.49 | 0.231 |  |
|  | pTimeShockZone | train | 1, 32 | 1.24 | 0.274 |  |
| Initial training (T1-T3) | NumEntrances | train | 1, 96 | 199.21 | 0.0 | *** |
|  |  | trial | 2, 96 | 17.47 | 0.0 | *** |
|  |  | train x trial | 2, 96 | 0.85 | 0.431 |  |
|  | Time1stEntr | train | 1, 96 | 16.21 | 0.0 | *** |
|  |  | trial | 2, 96 | 10.67 | 0.0 | *** |
|  |  | train x trial | 2, 96 | 8.27 | 0.0 | *** |
|  | pTimeShockZone | train | 1, 96 | 366.74 | 0.0 | *** |
|  |  | trial | 2, 96 | 2.98 | 0.055 |  |
|  |  | train x trial | 2, 96 | 1.29 | 0.279 |  |
| Initial recall (Rt) | NumEntrances | train | 1, 32 | 141.42 | 0.0 | *** |
|  | Time1stEntr | train | 1, 32 | 22.79 | 0.0 | *** |
|  | pTimeShockZone | train | 1, 32 | 411.4 | 0.0 | *** |
| Conflict training (T4-T6) | NumEntrances | train | 1, 96 | 36.77 | 0.0 | *** |
|  |  | trial | 2, 96 | 3.25 | 0.043 | * |
|  |  | train x trial | 2, 96 | 3.27 | 0.042 | * |
|  | Time1stEntr | train | 1, 96 | 27.9 | 0.0 | *** |
|  |  | trial | 2, 96 | 2.7 | 0.073 |  |
|  |  | train x trial | 2, 96 | 2.01 | 0.14 |  |
|  | pTimeShockZone | train | 1, 96 | 251.64 | 0.0 | *** |
|  |  | trial | 2, 96 | 0.62 | 0.542 |  |
|  |  | train x trial | 2, 96 | 2.12 | 0.125 |  |
| Conflict recall (Rn) | NumEntrances | train | 1, 32 | 49.7 | 0.0 | *** |
|  | Time1stEntr | train | 1, 32 | 16.36 | 0.0 | *** |
|  | pTimeShockZone | train | 1, 32 | 51.54 | 0.0 | *** |
| All trials (Pre-Rn) | PC1 ~ treatment | train | 1, 304 | 285.77 | 0.0 | *** |
|  | PC2 ~ treatment | train | 1, 304 | 28.61 | 0.0 | *** |
| Retention (Rn) | PC1 ~ treatment | train | 1, 32 | 46.74 | 0.0 | *** |
|  | PC2 ~ treatment | train | 1, 32 | 0.0 | 0.99 |  |

The conflict-trained animals show similarly rapid avoidance learning (T1-T3) as the standard-trained mice (Figure 3A-C). Initial between-day recall (Rt) of the conditioned place avoidance is also indistinguishable between the two Trained groups, and as expected, the Controls do not express an avoidance. Place avoidance in the two Trained groups differ when the shock zone was subsequently relocated for the conflict group (T4-T6), specifically on the first trial with the relocated shock location (T4) (Figure 3A-C). The conflict-trained group rapidly learns to avoid the relocated shock zone in subsequent trials, so that the ability to avoid shock cannot be distinguished between the two Trained groups (Table 1). Compared to the standard-trained group, the conflict-trained group learns to avoid two places in the same environment and extinguishes the avoidance of one location. Similarly, behavior is indistinguishable between the two yoked controls (Table 1). The day after the conditioning trials, on the memory retention test (Rn) with no shock, the conditioned avoidance persists and is indistinguishable between the two Trained groups, and is not expressed in the Controls, when tested the next day (Rn) under identical conditions for all mice.

Spatial memory is a complex phenomenon to which we aim to relate gene expression. In an effort to identify a memory score that does not privilege one estimate of conditioned avoidance over another we perform principal component analysis (PCA) on the 26 measures of behavioral space-use obtained from the automated behavioral analysis of position tracking (Table S1). Measures related to shock avoidance and avoidance conditioning are captured by PC1, which together reveals a “memory syndrome” that explains 37.4% of the variance when all the data from pretraining to retention are analyzed together. Following the experimental design, we focus the analysis on behavior during retention when the conditions were identical for all the mice, which was also just prior to euthanasia. At these times, PC1 explains 52.4% of the variance and PC2 explains 14.9% of the variance (Figure 3D). The variables with strongest loading on PC1 reflect avoidance behavior (Figure 3E), whereas PC2 captures measures of locomotor activity (Figure 3F), which can be considered an “activity syndrome.” PC1 significantly distinguishes Trained (n = 17) and Control (n = 17) mice (t_32_ = 6.8, p = 10^-8^), but PC2 does not (t_32_ = 0.01, p = 0.99). We designate PC1 as PC1_mem_ and use it as a comprehensive measure of the behavioral expression of active place avoidance memory, and PC2 is designated PC2_act_ (Table S1).

#### 2.2.2. Spatial memory drives hippocampal-subfield specific patterns of transcriptome variation

Transcriptomes were obtained from a subset of the mice (Figure S2). We begin by identifying differentially expressed genes (DEGs) between the four groups of trained and yoked mouse samples from the DG, CA3, and CA1 hippocampal subfields; 2 DG samples (1 standard-yoked and 1 conflict-yoked), 1 CA3 sample (1 conflict-trained) and 2 CA1 samples (1 standard-yoked and 1 conflict-yoked) are eliminated from analysis because of insufficient RNA reads. A total of 39 samples are analyzed (Figure S2). PCA distinguishes the trained and control DG transcriptomes, but not the CA3 and CA1 transcriptomes (Figure 4A). Few significant DEGs are found in any subfield between standard- and conflict-trained groups (Figure 4B,C) so we combine the data into one Trained (n = 9) and one Control (n = 9) group (Figure 4D). In the ∼17,000-gene transcriptomes we cannot detect any DEGs between Trained and Control CA3 samples, and only 4 in CA1 samples, whereas 263 DEGs are identified in the DG samples (Figure 4), with most (232) increased rather than decreased genes in Trained compared to Control samples (Figure 4D). We also tested the hypothesis that genes encoding the molecules implicated in the formation of LTP are associated with memory maintenance but of the 249 listed in ref (70), only 7 are in the DEG list (Table S3). The total DEG list in DG, CA3, and CA1 is available (see Table in Figure 4 legend).

**Figure 4.**
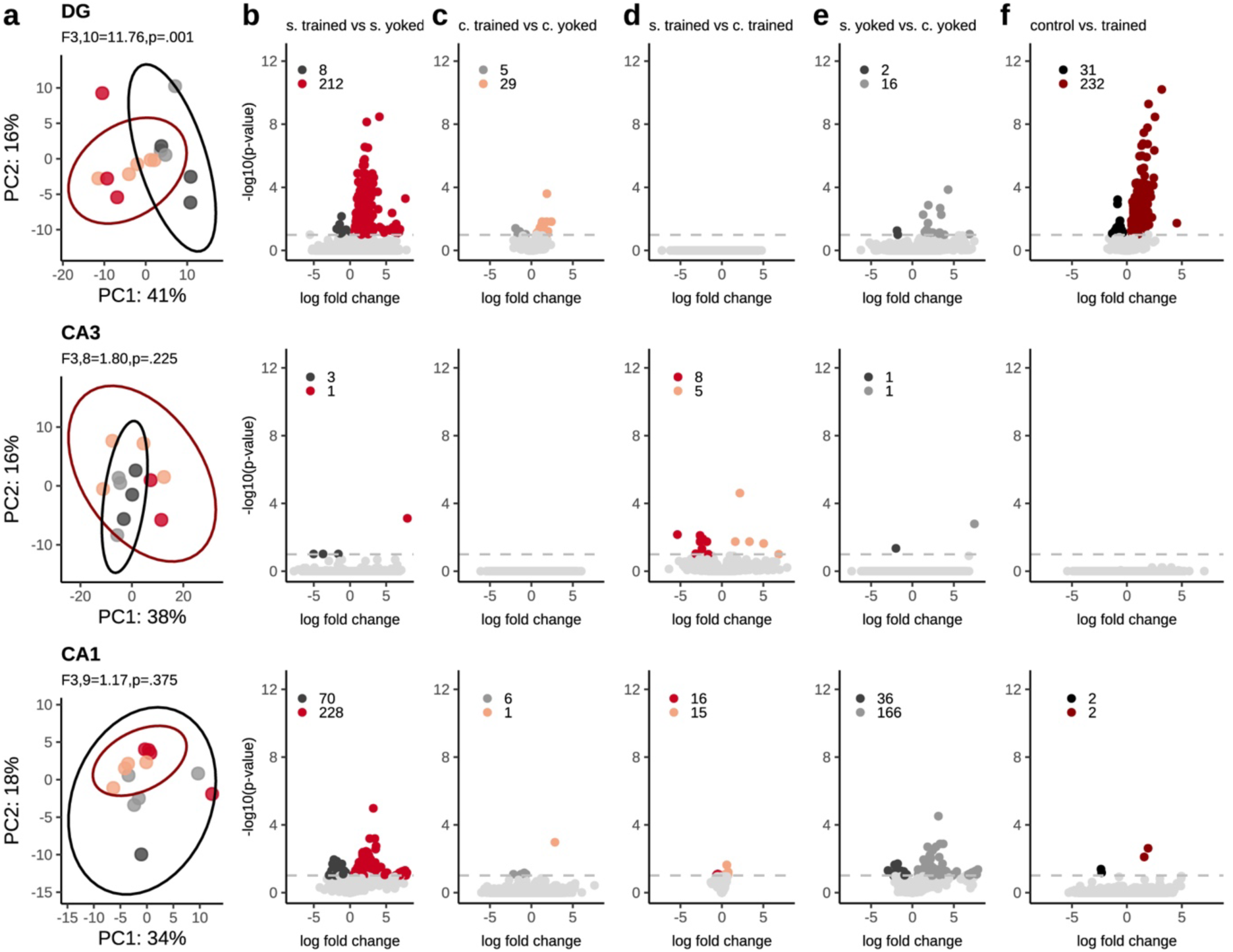
Subfield and treatment-specific gene expression differences. a) PCA can only partly distinguish trained and control DG transcriptomes. Volcano plots illustrating the significant and non-significant differences in gene expression when b) the standard trained and yoked groups, c) the conflict and yoked groups, d) the two trained groups, e) the two yoked groups, and f) the combined trained and yoked groups are compared. Rows show gene expression differences by region from DG where information enters the hippocampus, to CA3 and CA1, the output of the hippocampus. Each dot represents a gene whose color corresponds to whether its expression increased, decreased, or was not significantly different between the comparison pair. The total number of differentially expressed genes (DEGs) are reported in the top left corner of each panel. The log-fold change, adjusted p-value, log-transformed adjusted p-value, and directionality of significant gene expression in the DG, CA3, and CA1 are available here: <u>DEGlist</u>.

While *Prkcz* and related transcripts are not differentially expressed, the IEGs that are also DEGs are significantly correlated with PC1_mem_ (Figure 5A), and their pairwise correlations are positive (Figure 5B). These IEG relationships are not observed in CA3, which instead only identifies negative Pearson correlations between memory and *Prkci* that codes for PKC𝜄/λ and *Gria2*, the gene for the GluA2 AMPA receptor subunit (Figure 5C). *Gria2* is also significantly correlated with *Gria1,* the GluR1 AMPA receptor subunit, and *Grin1*, the NMDA receptor subunit GluN1 (Figure 5D). Furthermore, within CA3, the genes for all the candidate kinases forms a co-expression network (CaMKII, PKC*β*, PKMζ, and PKC𝜄/λ) with *Wwc1*, the gene for KIBRA/WWC1 that targets PKMζ to activated synapses (Figure 5D). In CA1, while expression of none of the candidate genes is linearly correlated with behavioral memory (Figure 5E), the pattern of significant pairwise correlations is intermediate to the DG and CA3 patterns, in part because *Gria1* expression covaries with the mutually correlated IEGs and also covaries with *Prkcb* and *Wwc1* (KIBRA), which itself correlates with *Prkcz,* the gene for PKMζ (Figure 5E).

**Figure 5.**
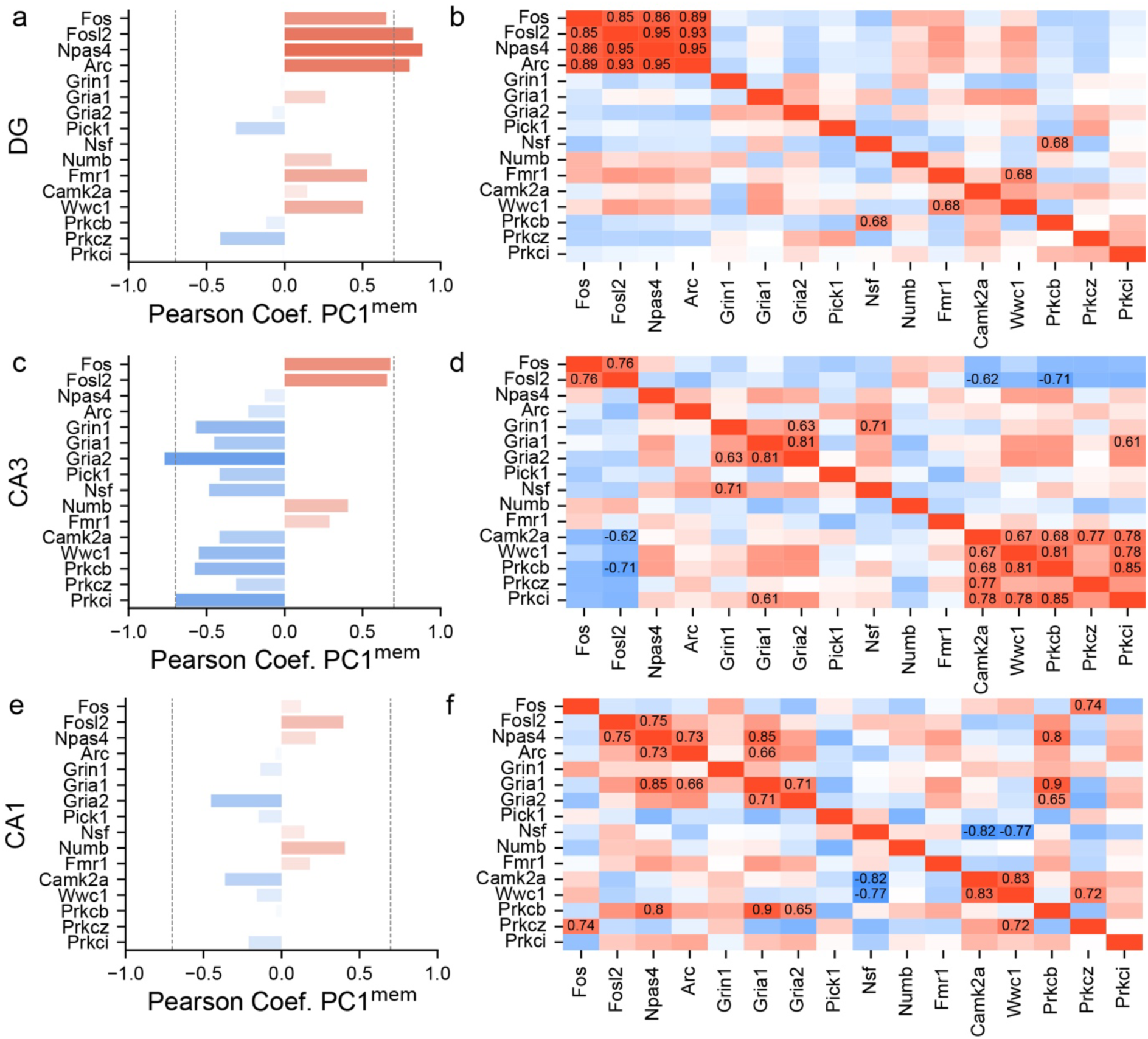
Hippocampal subfield-specific LTP and memory-associated candidate gene analysis. Pearson correlations of the expression levels between behavioral memory and candidate gene expression, as well as between pairs of candidate genes in a,b) DG, c,d) CA3, and e,f) CA1. Values are given for significant correlations (Absolute score ≥ 0.6). The correlation for the full set of analyzed genes is available here: <u>PC1_DEGCorrData</u>.

Astrocytes contribute to synaptic function and long-term memory persistence (71-74) and indeed, several astrocyte-associated genes are amongst the CA3 DEGs and some of their expression values in CA3 appear to covary with PC1_mem_ (Figure S3). While this is not the case for these genes in the DG or CA1 samples, nonetheless, many of the astrocytic genes are correlated with each other within the samples from DG, CA3, and CA1 (Figure S3).

The results of the preceding analyses vary substantially across the hippocampal subfields, so we examine whether the findings are robust or spurious. For one, the above analyses assume linear relationships between gene expression and memory or *bona fide* memory-related molecules like PKMζ, but some of these relationships are known to be non-linear due to regulatory, feedback and feedforward processes (11, 56). We therefore investigate whether associating these DEGs with memory is robust across three independent information-theoretic and statistical measures for ranking genes in the DG samples according to their ability to differentiate between Trained and Control median-of-ratio normalized gene expression levels. Indeed, Kullback-Liebler Divergence (KLD), mutual information (MI), and the Mann-Whitney tests all identify the IEGs and many of the DEGs identified by DESeq2, but not PKMζ (Figure S4). In particular, the LTP-associated genes *Arc*, *Egr1*, *Npas4*, and *Sgk1* all extremize these measures, indicating that they are most strongly associated with memory persistence.

#### 2.2.3. Might covarying gene co-expression networks predict shadow proteins better than individual genes?

Our observation that both the memory/LTP-related candidate genes, as well as the astrocytic candidate genes form co-expression networks indicates that instead of defining a high-dimensional state space of possible gene expression patterns, gene co-expression defines a lower dimensional state space. The gene-gene correlations define the intrinsic dimensions of the state space within which the Training and Control conditions are defined. We next ask whether such gene co-expression networks can reveal molecules in the “shadow proteome” such as PKMζ.

While the results of linear Pearson and non-linear Spearman correlations are largely consistent, both fail to detect non-linear but biologically meaningful relationships that can be measured by Chatterjee’s *ξ* (Fig. S5; 75). Since the DEGs distinguish Trained from Control samples, it is not surprising that 94% of the DEGs have significant Pearson correlations with PC1_mem_. We compare Pearson, Spearman, and Xi (*ξ*) measures of relatedness to PC1_mem_ (Figure 6). While the relationship of gene expression to PC1_mem_ for some genes is well characterized by the three measures, for some it is not. Because of its sensitivity to outliers, subsequent analyses use Spearman instead of Pearson correlations. In addition, to capture non-linear relationships like sigmoids and thresholds, we use *ξ* in a combined measure, Correlation Signal (Σ), defined as the norm of the two Spearman and Xi correlation measures (see Methods).

**Figure 6.**
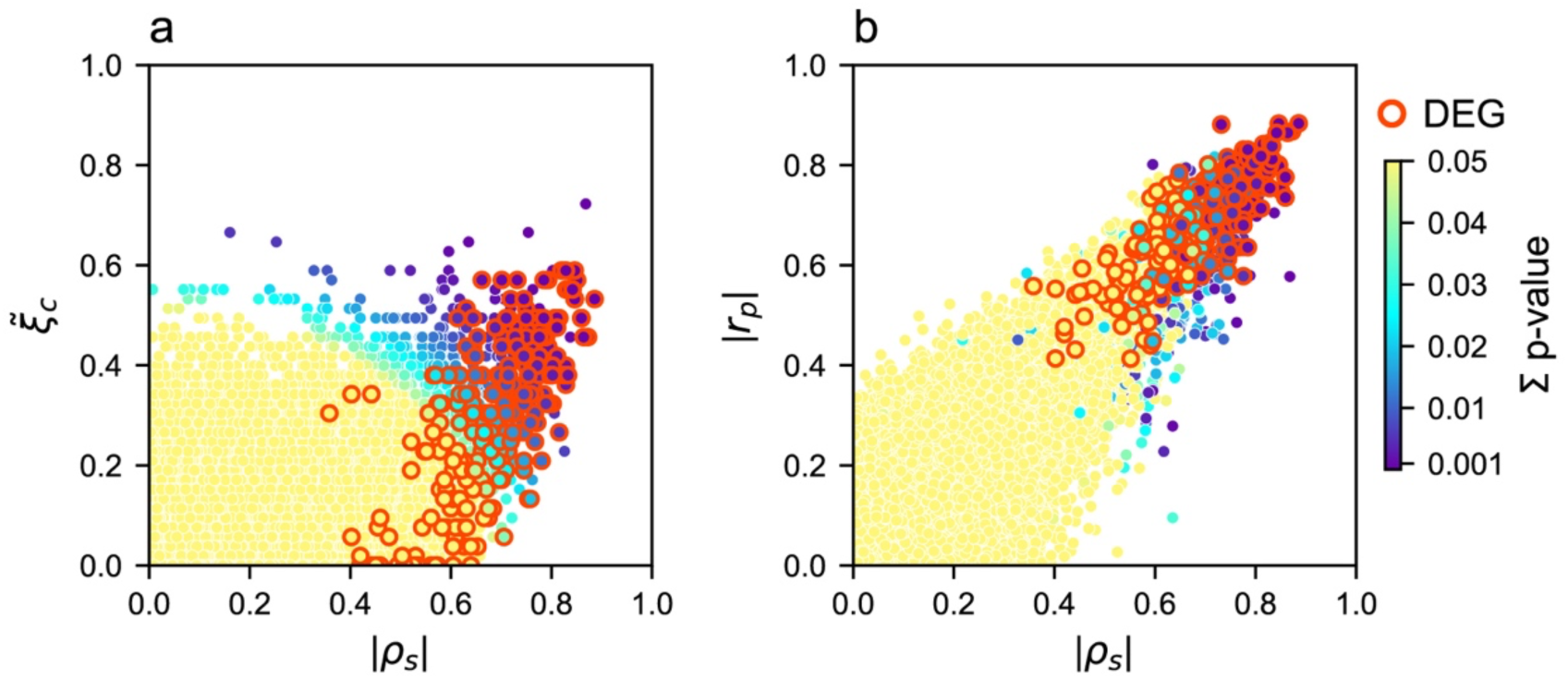
Comparing Pearson (*r*_*p*_, linear), and non-linear Spearman (*ρ*_*s*_), and Chatterjee *ξ̃_c_* correlations of gene expression and memory. The relationship between gene expression and PC1_mem_ was computed by the three correlation measures. Their significance (p ≤ 0.05) according to the Correlation Signal (Σ) of both *ξ* and ρ is color-coded. Comparison of a) Spearman and ξ correlations, and b) Spearman and Pearson correlations.

#### 2.2.4. Shadow proteins: differential gene expression does not identify PKMζ and related proteins in long-term memory persistence

Active place avoidance memory training increases PKMζ protein in specific hippocampal subfields (Figure 1). This complements findings from biochemical and genetic manipulations that *de novo* synthesis of PKMζ and its interactions with KIBRA and other proteins in hippocampus are crucial for the persistence of the active place avoidance memory (11, 16, 45), as well as other types of memory (3, 17, 20-23, 25, 26, 28, 29, 56, 57), even though genetic deletion of PKMζ is compensated by increased expression of PKC𝜄/λ (11, 76, 77). Nonetheless, candidate LTP/memory-persistence genes that differ significantly between the Trained and Control groups in the DG, CA3, or CA1 samples, are not related to the behavioral measure of memory (PC1_mem_), despite the memory-associated increases in the PKMζ protein with large effect size (Cohen’s d: DG = 1.3, CA3 = 1.5, CA1 = 2.5; power analysis with d = 1.5 predicts n’s of 7 are sufficient to detect RNA differences at α = 0.05 and *β* = 0.8). These data indicate that detection and discovery of such ‘shadow’ proteins is unlikely from standard transcriptomic analyses, leading us to consider innovations that can detect memory-related expression of PKMζ, and related molecules.

Weak-pairwise correlations amongst the many (i.e., high-dimensional) components of a system can indicate strong low-dimensional network states, in which single components are not crucial, but the collective interactions are strong determinants of function (78). Such low-dimensional manifold interactions are a familiar feature in studies of correlated neuronal action and field potentials (78-80) that have been proposed to replace the reliance on individual neurons (the neuron doctrine), with a population doctrine that relies on neuronal interactions to explain system functions (81). Accordingly, the positive identification of PKMζ as a shadow protein motivates exploring whether networks of transcriptomic interactions rather than single genes, may better predict the importance of proteins for memory persistence.

To test this idea, we define gene-gene networks (communities) of strongly correlated genes that are weakly correlated across communities. We first select 714 unique genes, comprising the 702 Behaviorally Relevant Genes (BRG) that are significantly correlated with PC1_mem_ at Correlation Signal p-value’s ≤ 0.05, as well as twelve additional candidate genes (CG). To infer gene co-expression networks that predict the behavioral Trained/Control groups, and PKMζ, we begin with the weighted gene co-expression network analysis (WGCNA; 82). Although WGCNA yields generally consistent modules when different correlation metrics are used, the assignments are unstably sensitive to algorithm parameters (Fig S6). To overcome this limitation, we use the Louvain Method of community detection (83). We construct an undirected graph G, in which each node represents a gene and edges represent the Correlation Signal, Σ, as the relationship metric between the 714 genes, and perform the Louvain clustering analysis 1,000 times with randomly selected seeds to assess community stability. We then examine whether this multiple-clustering approach reflects the underlying biological system by checking each gene’s co-cluster probability with pre-defined markers. Figure 7A shows the accumulated community assignments for the set of 714 genes. We find that these genes are reliably assigned to one of the three groups, each defined by the presence of the genes *Arc*, *Nsf*, and *Prkcz*, to which we assign red, green, and blue colors, respectively for visualization (57, 84, 85). The top-20 expressed genes in each of the three communities and their associated candidate genes reveal three robust gene networks (Figs. 7a, S7). Specifically, the IEGs are part of the red ensemble defined by *Arc* membership.

**Figure 7.**
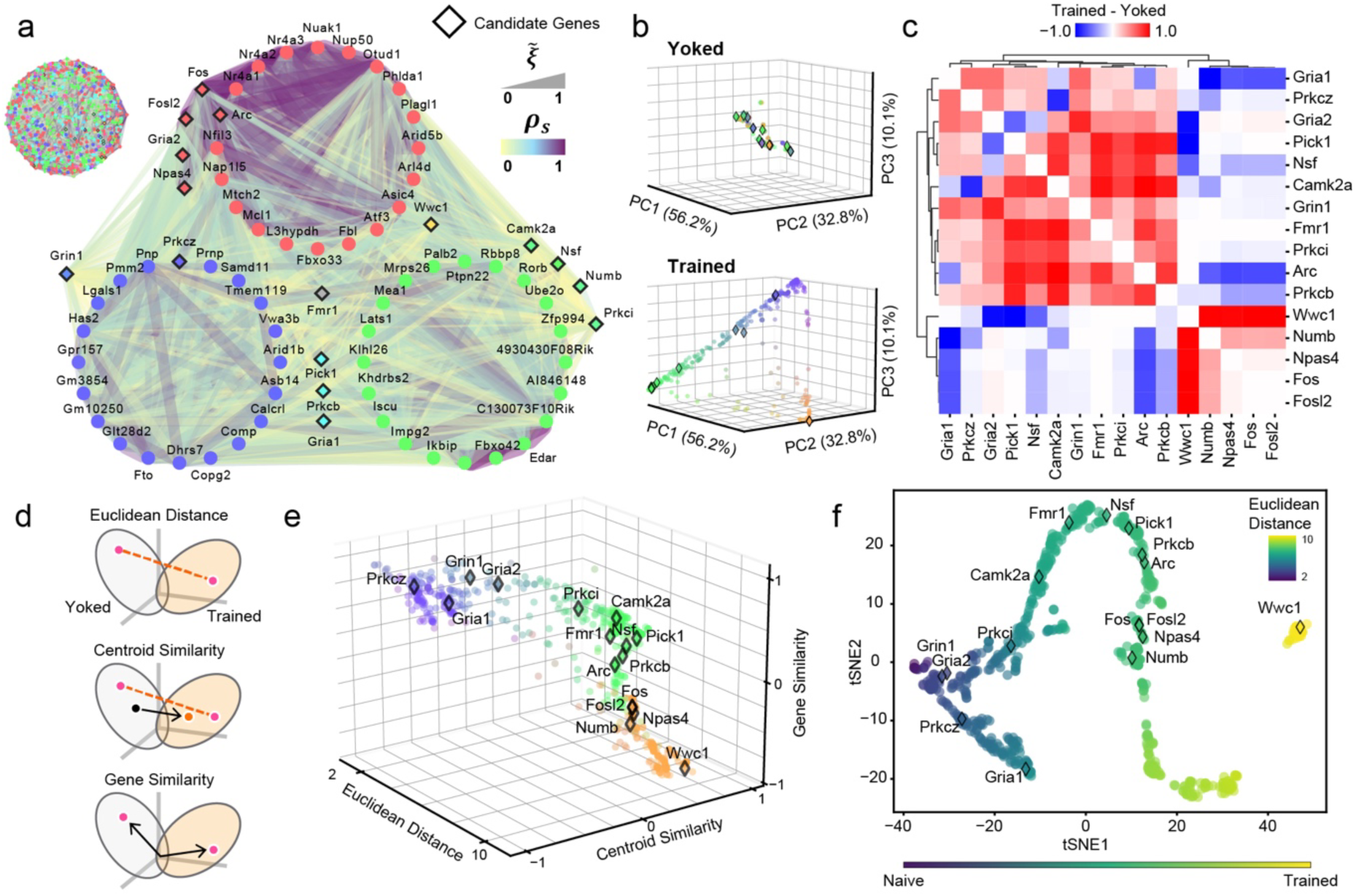
Co-Assign Probability changes based on Correlation Signal reveal a memory manifold. 714 genes were organized into communities by 1,000 iterations of the Louvain Method of community detection, each seeded with a different random set of genes. a) Validation of the new method through community co-assignment with three key genes. The community in which *Arc* is a member is always designated ‘red’, the community in which *Nsf* is a member is always designated ‘green,’ and the community in which *Prkcz* is a member is always designated ‘blue.’ Genes assigned to another community are designated ‘gray.’ After 1000 runs each gene has a red, green, and blue score with brightness set by the gray score. a - left) gene network with community color-coded genes and edges coded according to correlation values. a - right) The top 20 genes in each community and some candidate genes are organized according to their community color-code and the set of correlation values. The candidate genes are marked by the diamond outline. b) Separate yoked and trained 714-D Co-Assign Probability (CAP) matrices were computed to estimate the probability of all the gene – gene co-clustering probabilities. The two clustering probability matrices were then projected into a common 3-D PCA space (ℳ), generating *<u>r</u>_<u>y</u>o_* and *<u>r</u>_<u>t</u>r_*. c) Hierarchical clustering of the difference between the trained and yoked CAP matrices, using Pearson correlations as the distance metric. d) Cartoons illustrate calculations of the Euclidean Distance, Centroid Similarity, and the Gene Similarity measures of how the yoked and trained genes changed within each condition’s projection in ℳ. e) Summary plot of how the 714 genes are distributed across the features depicted in panel d. Colors in panels b and e correspond to their positions within the value distributions. f) t-SNE plot derived from the three metrics for dimensionality reduction. The color bar represents the magnitude of Euclidean Distance changes from the yoked to the trained data set.

Genes associated with early-LTP and the trafficking and insertion of AMPA receptors associate within the green ensemble defined by *Nsf* membership. Finally, the NMDA receptor gene *Grin1* is part of the blue ensemble defined by *Prkcz* membership. Note that *Wwc1* (KIBRA) is uniquely yellow, indicating that it is assigned with similar probability to the red ensemble enriched with IEGs, and the green ensemble enriched with genes associated with early-LTP and AMPA receptor placement. Genes associated with PKMζ-mediated effects on AMPA receptors, *Pick1, Gria1, and Prkcb* are identified as turquoise because they are reliably assigned to either the green ensemble or the blue ensemble. It is also remarkable that *Fmr1* is gray as it is assigned approximately equally to each of the three communities. This suggests there may be a detectable functional logic to the organization of these gene ensembles.

To further validate this network approach, for each iteration, we record how often each pair of genes clusters together, building a 714 × 714 gene-gene Co-Assign Probability (CAP) matrix. We apply PCA to this matrix to project genes into the reduced 3D CAP space (ℳ) defined by the top three principal components and obtain the vector of gene coordinates (<u>*r*</u>). To assess whether clustering patterns change with memory training, we compute CAP matrices separately for the yoked and trained conditions. These matrices are then projected into the same PCA space (ℳ) as the total dataset to ensure direct compatibility (Figure 7B), yielding <u>*r*_*y*_</u>_*o*_ and <u>*r*</u>_*<u>t</u>r*_ for the yoked and trained groups, respectively. To enhance visual contrast, we normalize PCA coordinates and convert them to hex color codes. By inspection, yoked gene-gene relationships appear compact, while trained relationships are more dispersed along triangular axes (Figure 7B).

For comparison, we also perform PCA directly to the raw gene-expression matrix (genes × samples) after z-transforming each gene’s expression across all samples. PCA of the raw-expression values does not clearly separate the candidate genes or distinguish yoked from trained conditions (Figure S8). PCA performed on the raw Correlation Signal (Σ) and the WGCNA Topological Overlap Matrix (TOM) similarity scores (Figure S9) produce generally spherical distributions, indicating that while they can capture meaningful biological relationships to some extent, their spherical structures lack clear separation. In contrast, PCA performed on the Louvain clustering-based CAP matrices distinctly highlights differences between conditions and gene groups, underscoring the advantage of explicitly examining gene-gene relationships using clustering probabilities.

To further understand how co-clustering patterns differ between training and yoked conditions, we generate hierarchical cluster using the separate yoked and trained CAP matrices (Figure 7C). We first subtract the yoked clustering probability matrix from the trained matrix and perform hierarchical clustering using Pearson correlation as the distance metric. The range of the heatmap is -1 to 1, which corresponds to losing and gaining co-cluster likelihood, respectively. In the heatmap of all 714 genes, *Prkcz* groups with *Nsf* but not with *Numb*, consistent with known cellular roles in synaptic plasticity, where PKMζ and NSF promote potentiation and NUMB promotes depotentiation. Additionally, *Prkcz* is grouped together with the genes associated with the endoplasmic reticulum related gene ontology terms.

To highlight the genes that are driving the differences between the yoked and trained conditions, we calculate a shift vector (Δ*C* = *<u>r</u>_<u>t</u>r_* - *<u>r</u>_<u>y</u>o_*) in PCA space (ℳ) for each gene. From ΔC, we derive: (1) the Euclidean Distance (∥ ΔC ∥) — overall magnitude of each gene’s shift; and (2) the Centroid Similarity 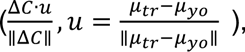 which is the cosine similarity between each gene’s shift and the global centroid shift between the yoked and trained conditions; and (3) the Gene Similarity 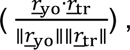 which measures the cosine similarity between the yoked and trained conditions for each gene (Figure 7D). We observe that the genes distribute along a low-dimensional manifold characterized by these three measures (Figure 7E), which we confirm by t-SNE non-linear dimensionality reduction (Figure 7F), and by visualizing all pairwise 2D combinations of the three features (Figure S10). The distribution along the manifold reflects the Euclidean distance from the yoked condition. Notably, the KIBRA gene *Wwc1* is at the extreme of the manifold, distinct from the other candidate genes that are separately distributed along the manifold. We also validate this result using another dimensionality-reduction technique, Uniform Manifold Approximation and Projection (UMAP) (Figure S11). Even when we select alternative gene-gene scores such as the absolute value of the Spearman or Xi correlations instead of their norm, the Correlation Signal (Σ), the overall result is robust, highlighting that the change in co-clustering probability is the most effective factor in the search for important genes that contribute to maintaining memory, despite being hidden in the shadows of the most abundantly expressed genes (Figure S12).

## 3. Discussion

### 3.1. Summary and Limitations

We graphically represent our overall approach in Figure S13. To detect changes in the expression of genes for PKMζ and other proteins known to be important for long-term memory, we first assessed gene expression changes at the individual-gene level by identifying differentially expressed genes, few of which corresponded to proteins known to be important for long-term memory (Figure 4,5). Next, we examined gene expression shifts at the population level, considering all genes significantly altered by training (Figure 6) and a set of candidate genes hypothesized to respond to memory training by observation or their biochemical relationship with PKMζ. To capture these population-level changes, we clustered genes according to their Correlation Signal across the two conditions rather than relying on individual gene-wise differential expression. For each gene, we then estimated its cluster co-assign probability and compared how those probabilities differed between conditions. Overall, our results demonstrate that active place avoidance memory training alters the covariance structure among individual gene expression changes (Figure 7).

We began by evaluating long-term memory for the conditioned avoidance response in active place avoidance memory. We first measured that PKMζ protein is persistently increased in the somatodendritic compartments of the trisynaptic pathway for at least a month (Figure 1). We then quantified both recall of the conditioned behavior and the associated transcriptional changes in three subfields of the hippocampus a day after the end of the training and 30-min after expression of the avoidance memory. The behavioral similarities and differences between the Trained and Control groups reflect knowledge about the location of shock rather than the experience of shock *per se*, because the 24 hours before euthanasia were physically identical across the groups, constituting time in the home cage and 10 minutes on the arena with no shock, and the comparison groups experienced the identical physical conditions. We cannot assert that the gene expression findings associated with active place avoidance memory are due to altered gene expression specifically in neurons because we sampled bulk tissue, containing approximately equal numbers of neurons and other cells that are largely the glial population (86).

We started with an unbiased data-driven approach to identify genes that are sensitive to the maintenance of spatial memories across at least 24 h, then conducted a hypothesis-driven analysis of the data to characterize whether the memory training is associated with altered expression of dozens of genes that have been previously shown to be important for learning, memory, and the cellular model LTP, including PKMζ that we confirmed is increased two-fold at each of the subfields we sampled for transcriptional profiling. Because prior work showed that memory-related PKMζ originates from *de novo* synthesis within 2.5 h of active place avoidance training and PKMζ is increased 24 h afterwards in trained but not yoked control mice (11, 15, 66), we expected to find transcriptional evidence of memory-related PKMζ gene expression, unless it is not transcriptionally regulated. For this hypothesis-driven approach, we examined both differential gene expression and its covariance with estimates of avoidance memory behavior and with other candidate genes. Our RNA-seq methods could not detect ribosomal RNA, nor did we perform protein sequencing, limiting the scope of this study to findings that may related to memory-related transcriptional regulation.

### 3.2. The PKMζ engram

Richard Semon introduced the term “engram” to mean the physical trace that stores memory, which is a concept that Plato introduced by proposing memory is like a block of wax in which perceptions and thoughts are imprinted, “just as we make impressions from seal rings” (87, 88). We observed that PKMζ is persistently increased for at least 1 month, selectively in memory-tagged cells and specifically at synaptic populations of the trisynaptic pathway (Figure 1). We did not demonstrate if this PKMζ trace is necessary or sufficient for maintaining the conditioned place avoidance behavior; however, prior work discovered place avoidance training potentiates the Schaffer collateral synaptic population at CA1 *str. radiatum* but not at CA1 *str. lacunosum moleculare* (10) and at the middle molecular layer of supraDG, complementing our current findings (Fig. 1; refs 5, 66). Notice that the compartment-specific persistent increases of PKMζ in DG, CA3, and CA1 point to an active process targeting the protein, since PKMζ is observed throughout the hippocampus circuit, but is elevated selectively by memory training. The compartmentalization of the PKMζ increases that we observe is hard to reconcile with a somatic synthesis of PKMζ and its free diffusion in cytosol to activated synapses. However, compartmentalization is compatible with local translation and recent findings that PKMζ complexes with the cytoskeleton-associated postsynaptic scaffolding protein KIBRA that persistently targets the kinase’s action to activated synapses to store memory (45, 89). These findings together suggest the PKMζ trace is a *bona fide* engram because i) PKMζ is both necessary and sufficient for maintaining late-LTP in wild-type mice, and active place avoidance memory in wild-type mice is eliminated by ii) inhibition of PKMζ synthesis during learning, and, for up to a month after the avoidance memory is established, it is erased by iii) inhibition of PKMζ kinase activity or iv) inhibition of PKMζ binding to KIBRA (11, 16, 21, 45, 90). Systems consolidation, taken at face value, might suggest that memory no longer depends on hippocampus after a month, but that is falsified by the cited work as well as demonstration of a core and persistent hippocampal dependence of other forms of memory, despite the possibility of systems consolidation (91-94). Accordingly, the inability to detect simple transcriptomic differences in LTP- and memory-associated genes cannot be attributed only to systems consolidation at the early time points after training that we examined. Our observations of the PKMζ trace are consistent with it being a memory trace by definition, and arguably a long-pursued engram.

### 3.3. Memory-associated Differentially Expressed Genes

Gene expression differences between trained and control groups are mostly observed in DG, with substantially more increases of gene expression than decreases in the trained mice compared to the yoked controls. Few genes were differentially expressed in Ammon’s horn (Figure 4), which is not expected in light of theories of hippocampus memory and information processing (36, 37, 95) and findings that maintenance of the active place avoidance memory persistently enhances CA3-CA1 synaptic function for at least 30-days (10). Nor did we find memory maintenance-related differential expression of genes for the molecular translation machinery in the MEK/ERK and MTOR pathways, important for memory-related transcription and translation during initial cellular memory consolidation (96, 97). Several genes that are differentially expressed in DG and are positively correlated with avoidance behavior are IEGs, including *Arc* and the transcription factors *Fos*, *Jun*, and *Npas4*. In addition to being differentially expressed, expression of these genes is highly correlated with one another. In previous experiments, *Arc* was activated during contextual threat avoidance conditioning and optogenetically silencing the *Arc*-activated DG neurons was sufficient to impair contextual fear memory, whereas optogenetically stimulating neurons that activated *Fos* was sufficient for expressing the memory (98). Indeed, as we also observed in DG, prior work showed that *Arc* and *Fos* expression are correlated in active CA1 neurons (99). Because we examined gene expression a day after training when training-associated IEG activation is not expected (100, 101), it is possible that the robust IEG activation we observed is related to the mice learning about the absence of shock during the retention test. However, this explanation predicts an inverse relationship between PC1_mem_ and IEG expression, because mice that expressed the weakest place avoidance by entering the shock zone the most during the retention test (low PC1_mem_ values) would have the greatest opportunity to learn that shock was absent and the five mice that showed perfect avoidance by never entering during the retention trial could not have learned that the shock was off. We reject this alternative because the correlations between IEG expression in DG and PC1_mem_ on the retention test are all positive. Instead, these findings suggest that IEGs can respond not only to the novel experience of stimuli that are encoded during the conditioning of a response, but that their activation in DG also corresponds to the expression of long-term memory (102, 103).

Gene ontology analysis revealed that many of the genes that are differentially expressed in the DG exert their function in dendrites and are important for neuronal projections and cell-cell junctions, including synapses, which is expected from the relationship between LTP and memory that forms the basis of the widely-accepted synaptic plasticity and memory hypothesis (Table S2; 33). Although, we cannot distinguish the contributions of neurons and glia to the memory-related differential gene expression we observed, these findings suggest that the expression of the conditioned place avoidance altered neural activity in a way that increases transcriptional activity, perhaps for the purpose of promoting dendritic growth. Indeed, we note that expression of a number of astrocyte-related genes, especially in CA3 were positively correlated with the PC1_mem_ estimate of memory expression, including *Aldh1a1*, *Aldh1l2*, *Gfap*, *Fgfr3*, and *Aqp4* (Figure S3). Even though these genes were not differentially expressed after avoidance training, the correlation with expression of the conditioned avoidance behavior adds to the growing evidence that astrocytes also play an active role in forming and/or stabilizing memory via a variety of potential mechanisms (74, 104-107).

Synaptic reorganization requires that neurons polarize their dendrites to create new neural connections and direct channels to specific regions within the synapse. Disruption of the processes that regulate synaptic organization can impair memory or cause disease. Dysregulation of *Eif5* a DEG, is associated with many neurodegenerative diseases (108). *Cpeb4*, a DEG, encodes cytoplasmic polyadenylation element binding protein 4, which regulates the translation of genes by modulating their poly(A)-tail length and is a known regulator of genes associated with autism spectrum disorder (109). Many of the genes that are mutated in autism spectrum disorder are crucial components of the activity-dependent signaling networks that regulate synapse development and plasticity (110). Deletion of *Fmr1* can lead to dysregulated mRNA translation, altered synaptic function, abnormal protein synthesis-dependent synaptic plasticity, and poorly coordinated neural discharge (61-63). Very few of the differentially expressed genes in our study overlapped with the LTP-related molecules that were identified from review of the LTP literature (Table S3; 70). Some classes of genes that are notably absent from the list of differentially expressed genes but present in the LTP-related list are genes for receptors (calcium, glutamate, GABA, and serotonin), calcium or calmodulin-binding proteins, and adhesion molecules. This is perhaps not surprising because the review focused on literature examining LTP induction, the first minutes of long-term potentiation, whereas our experiments were designed to investigate 24-h memory expression, which would be expected to correspond to LTP maintenance. Thus, this work is consistent with the evidence (Table S3; reviewed in 57) that like LTP-induction, many molecules known to be crucial for learning, are neither crucial nor expressed in LTP maintenance and as we observe, memory persistence. This points to different molecular mechanisms for forming a memory and maintaining and/or later expressing it days later.

### 3.4. Transcriptional profiling did not identify genes known to encode proteins crucial for LTP and memory persistence

What then of the atypical PKCs and CaMKII, candidate molecules that are demonstrated to have crucial roles in the maintenance of LTP and synaptic structure, as well as active place avoidance memory, as demonstrated in Figure 1 (11-13, 15, 16, 29, 45, 66)? We did not detect differential expression of these genes, nor did we detect correlations between any of the corresponding mRNAs and the behavioral measures of avoidance memory expression, whereas we did for IEGs as discussed above. It would, however, be naive and/or hasty to conclude that despite being unbiased, this transcriptomic screen contradicts the findings of the candidate molecule approaches that have identified a role for these molecules, especially in active place avoidance memory. This highlights a major shortcoming of using transcriptional profiling to identify molecular mechanisms of functions such as memory. First, the abundance of mRNA does not relate in a straightforward way to the cellular abundance of protein because the relationship depends on the translational regulation of the particular protein as well as protein translocation and stability (111, 112). Indeed, as an example, while PKMζ is strongly regulated by repressing translation at dendrites (113), there is no evidence that it is transcriptionally regulated, either in late-LTP (54) or in active place avoidance conditioning (15), in line with the present findings. While this highlights a limitation of transcriptional profiling, it draws attention to the necessity of performing combined transcriptional and proteomic profiling in tissue taken from the same subjects (112). In light of the present findings, the hypothesis that PKMζ’s molecular interactions maintain memory (13, 45, 56, 114) predicts independence between *Prkcz* RNA that encodes PKMζ, and measures of memory in an unbiased screen that integrates memory-related measures of behavior with circuit-specific transcriptional profiling (RNA), as we have done and observed here. Somewhat unintuitively, the hypothesis also predicts that in the same tissue, memory measures correlate positively with PKMζ protein levels and those of its interaction partners (e.g., KIBRA, NSF, NUMB, PICK1, GluA2), even though their RNA levels show little phenotypic variation (15, 48, 50, 55, 85, 115), similar to observations in LTP (116). There is likely substantial utility in such analyses that use experimental design and concurrent multi-scale measurements from single subjects to integrate information from across levels of biology (here behavior, protein and gene expression), not only to discover unknown and unanticipated mechanisms of complex, poorly defined processes like memory, but to also test the validity of specific mechanistic hypotheses. For example, the notion that the atypical PKCs may play complementary but competing roles in the maintenance of LTP and memory, perhaps through binding to KIBRA, predicts anti-correlation between *Prkcz* and *Prkci* and whichever isoform is more expressed, to be correlated with KIBRA. This pattern was however not observed, suggesting a simple model of transcriptional regulation does not map their functional protein-mediated functional relationships. Although we have observed it is not straightforward, the findings nevertheless suggest that the transcriptional level of biology can relate to biological function that is mediated by proteins, even rare ones like PKMζ.

### 3.5. Illuminating the shadow proteome

Taking a lesson from a major conceptual advance in analysis of neuronal spike train populations, we also considered whether hypothesizing straightforward relationships between gene expression (or even protein expression in principle) is conceptually fraught, in part because individual expression changes can have manifold interactions with network-level consequences (81, 117). We first, demonstrated that non-linear measures of correlation like Chatterjee’s *ξ* identify *bona fide* relationships that the standard use of Pearson’s correlation failed to identify (Figure 6). We then used the combination of Spearman’s ρ and ξ correlations to define ensembles of mutually correlated genes (Figs. 5,7). Initial skepticism was assuaged because ensembles of mutually correlated genes were reliably identified, i.e., their membership was meaningful according to known biochemical function, and perhaps most significantly, this approach identified candidate genes independently proven to be involved in memory, even in area CA3 where a co-expression network was observed amongst CaMKIIα, PKC*β*, PKMζ, PKC𝜄/λ and *Wwc1*, even though no DEGs were detected between trained and control samples (Figs. 4F, 5D). This last fact suggests it may be possible to identify both known (as shown here) and unknown components of the shadow proteome by challenging the conventional hypothesis that there are unique, essential components for maintaining memory. Put simply, function emerges from the subtle, cooperative tuning of many components, and not from a dramatic change in any one, just as an arch bears a bridge’s weight through the balanced interplay of all its stones, not by any single one. Consider for example, the ongoing debate as to whether the persistent kinase CaMKII or PKMζ is crucial for persistent memory storage, or whether some other molecule is crucial, at the exclusion of all others (118-120). Neither kinase, nor the gene for the compensating kinase *Prkci* were found to be differentially regulated between the memory and control groups, nor did memory measures correlate with the expression of these kinase genes or with the expression of their scaffolding partners that guide their function, like KIBRA. However, by abstracting both linear and non-linear gene-gene relationships to define the concept of a gene Co-Assign Probability network, such distinct gene ensembles could be i) reliably defined, and ii) were the dataset’s best predictors of the involvement of genes that encode shadow components, undetected but known proteins that are crucial for the memory being investigated (Figs. 1,7).

## 4. Conclusion

Our findings suggest a hypothesis, which contends that a low-dimensional manifold of molecular interactions (genes, mRNAs, proteins) is embedded in the high-dimensional state space of biological components. The manifold dimensions are defined by biological interactions/functions, not by the biological components themselves. This view extends the manifold concept from electrophysiological studies to other levels of biology, where the high-dimensional space of potential memory-maintaining biological components, in practice, collapses into a low-dimensional manifold of biological solutions for maintaining memory (79, 80). In this view, the logic of the functional implementation, rather than the traditional, parallel measurement of physical components is best able to identify the solutions. Despite vast heterogeneity in biological elements (genes, proteins, cell types with their own cell biology and electrophysiology that contribute to memory), this view asserts there is a conserved computational logic to how these elements interact to realize memory storage and expression. Innovative research programs will be required to validate or refute this conjecture (121), with important implications if it is supported. For example, if key biological functions like memory are governed by a manifold of multiple replaceable interacting genes and/or proteins, then the endeavor to find definitive “causal” biomarkers of function and dysfunction is at best fraught, because key components can be replaced when they are absent or defective (64, 122), as is the case when PKMζ is genetically deleted and is certainly the case for neuronal circuit functions and the cell-specific molecular phenotyping that has been compellingly demonstrated with the crustacean stomatogastric ganglion (STG) model system (11, 45, 123). On the other hand, as the STG as an archetype demonstrates, neuronal systems are robust and treatment options can be plentiful when the system components are manifold, replaceable, and can be coordinated (124-126).

## 5. Methods

### Experimental Subjects

All animal care and use complied with the Public Health Service Policy on Humane Care and Use of Laboratory Animals and were approved by the New York University Animal Welfare Committee and the Marine Biological Laboratory IACUC. Transgenic ArcCreER^T2^ (originally purchased from The Jackson Laboratories, Jax: 021881) and R26R-STOP^flx^-ChR2-EYFP mice (a gift from Christine Denny) were crossed to generate the ArcCreER^T2^-ChR2-EYFP mice for Experiment 1, as previously described (66, 127, 128). Wildtype C57BL/6J mice (originally purchased from The Jackson Laboratories, Jax: 000664) were used for Experiment 2.

#### Experiment 1 – Tracing Memory

Memory-induced increases of protein expression were studied to identify changes in the strengthening of hippocampal synaptic populations that persist for a month after memory training. We focused on PKMζ because it is both necessary and sufficient for the maintenance of LTP in hippocampal slices over at least five hours, and PKMζ in hippocampus is necessary for active place avoidance memory and its memory-initiated protein expression is enhanced in CA1 for at least a month (11, 15, 16, 66). PKMζ protein increases in synapses during memory storage, making this an ideal method to investigate the subcellular distribution of the protein within the different regions of the hippocampus (66). PKMζ visualization through immunohistochemistry (IHC) was combined with Targeted Recombination in Active Populations (TRAP2) technology to identify neurons that express a fluorescent marker (EYFP) once the expression is initiated by the external ligand, 4-hydroxytamoxifen (TAM, Sigma, St. Louis, MO, H7904). The Cre-mediated recombination is directed to neurons expressing high levels of the immediate early gene (IEG) *Arc* while TAM is circulating. The combination of these two techniques allows us after a month’s delay to identify the EYFP-expressing neurons as likely to have been active during TAM administration, and to interpret the synaptic sites within these neurons that exhibit elevated PKMζ levels as loci of persistent potentiation.

Thus, the ArcCreER^T2^ x ChR2-EYFP TRAP2 mice (n=4 Trained and n=4 Untrained) were temporarily housed in an experimental room with an isolation chamber (1 mouse/cage) on a 12:12 light/dark cycle. These mice contained a ArcCreER^T2^ DNA sequence allowing initiation of transcription at the *Arc* promoter to express Cre recombinase in the presence of TAM. The mice also contained a STOP codon flanked by LoxP sites followed by the sequence for Channelrhodopsin 2 (ChR2) fused with EYFP (Figure 1A). Thus, TAM will induce activation of Cre recombinase in *Arc* activated cells to remove the STOP codon to cause perpetual membrane-targeted EYFP expression. These ArcCreER^T2^ x ChR2-EYFP mice allowed us to tag neurons with EYFP if they activate *Arc* during expression of the conditioned place avoidance, as in prior work (66, 128).

To reduce spurious activation of *Arc* and EYFP tagging, 2-3 days before the initial exposure to the behavioral arena, all mice began housing in a sound-insulated and light-controlled isolation chamber that was approximately 1 m away from the behavioral apparatus. The mice remained housed in the isolation chamber for two days after TAM administration and afterwards were returned to the vivarium. This isolation limited the association of EYFP expression to the arena experience, as was validated in prior work (66). Thirty minutes before the third training trial when the mice expressed strong place avoidance memory (see below), all mice were injected with TAM (2 mg in 200 µl i.p.).

#### Experiment 2 – Transcriptional Profiling of Memory

The subjects for transcriptional profiling were a subset of the 34 C57BL/6J mice that went through the behavioral training protocol. They were housed at the Marine Biological Laboratory on a 12:12 (light: dark) cycle with continuous access to food and water in home cages with up to five littermates. Only male mice were analyzed in this study to limit gene expression variations due to estrus status.

### The Active Place Avoidance Task

We used an active place avoidance paradigm to establish a hippocampal-, synaptic plasticity- and PKMζ-dependent spatial memory that persists for months (5-7, 10, 11, 16, 45). Briefly, an overhead digital video tracking system monitors the position of a mouse as it moves on an elevated circular 40-cm diameter rotating (1 rpm) arena; the transparent arena wall allows the mouse to observe the environment (Tracker, Bio-Signal Group Corp., Acton, MA). The computer activates a mild constant current (0.2 mA, 500 ms, 60 Hz) foot shock across the parallel bars of the floor whenever the mouse enters a 60° stationary sector that is designated to be the shock zone. Mice rapidly learn to avoid the shock zone. Behavior in the arena is automatically quantified by 48 measures during each trial, as subset of which are used for formal analysis (Table S1; Tracker, Bio-Signal Group Corp., Acton, MA).

#### Experiment 1 – Tracing Memory

Mice were removed from the isolation chamber and placed on the behavioral arena and were subsequently returned to the chamber in a way that minimized novel or distinctive extra-arena experience. Each mouse received a 30-min pretraining trial (Pretrain) with no shock. After a 24-h interval back in isolation, each mouse received three 30-min training trials (Train 1-3), separated by 24-h intervals in isolation. Four Trained mice received memory training to avoid the location of a mild foot shock in a 60° shock zone; these mice acquired a conditioned place avoidance. Four other Untrained control mice received the identical experience except shock was never turned on. Thirty minutes before Train 3, each animal was injected with TAM. This was intended to tag conditioned avoidance memory-activated cells with ChR2-EYFP in the Trained group and control-activated cells in the Untrained group. All mice were returned to their home cage for 30 days before testing retention of the conditioned avoidance without shock.

#### Experiment 2 – Transcriptional Profiling of Memory

We used a three-day training protocol (Figs. 2a, S2). On day 1, the mice (n = 34) were acclimated to the rotating arena during a 10-min pretraining trial (P) with no shock. Each Trained mouse (n = 17) mouse then receives three 10-min training trials (T1-T3), each separated by 2-h intervals, after which they are returned to their home cage. The next day, the mice are retested (Rt), with the shock in the same location. This was followed by three additional 10-min training trials (T4-T6), each separated by 2-h intervals. The shock zone remained in the same location during T4-T6 for standard-trained animals (n = 8) but it was relocated 180^°^ for the conflict-trained mice (n = 9). The relocated position of shock coincides with the place the mice had preferred to be during the T1-T3 trials (Figure 2A). Each trained mouse (n’s = 8, 9) was yoked to a Control mouse (n’s = 8, 9) that received the identical shock time series (Figure 2B). Accordingly, Trained (n = 17) and Control (n = 17) mice had the identical physical experiences of the environment, but the Trained mice could learn to avoid the shock zone, and the Controls could not. On Day 3 all mice received a memory retention trial (Rn) with no shock to evaluate the conditioned avoidance. The Trained and Control mice were each divided into two groups that only differed during the T4-T6 trials on Day 2. Littermates were randomly assigned to one of the 2 (Standard-Trained/Conflict-Trained) x 2 (Standard-Yoked/Conflict-Yoked) groups (Figure 2A). Note that the shock is mild and compared to mice on the arena but never shocked, we do not detect behavioral or plasma corticosterone expression differences in these groups 24-h after the avoidance or yoked experience compared to the untrained of walking on the rotating arena (129).

### Immunohistochemistry and confocal microscopy (Experiment 1)

Detailed methods have been reported and are only briefly summarized here. Immediately after testing retention, animals were anesthetized with 2% isoflurane for two minutes, and the brains were fixed by cardiac perfusion then removed. Transverse 300-μm hippocampal slices were placed in ice-cold 4% paraformaldehyde in 0.1 M phosphate buffer (PB, pH 7.4) and post-fixed for 48 h, as previously described (66). Antibodies specific to EYFP (Abcam, #ab13970) and PKMζ (C2 polyclonal antiserum from the Sacktor lab) were used. Slices were then washed with PBS (pH 7.4) and 30-µm sections of the dorsal hippocampus were prepared using a Leica VT 1200S vibratome (Leica Biosystems, Buffalo Grove, IL). Free-floating sections were permeabilized with PBS containing 0.1% Tween20 (PBS-T) for 1 h at room temperature and blocked with blocking buffer (BB; 10% normal goat serum in PBS-T) for 2.5 h at room temperature. The sections were then incubated overnight at 4°C with primary antibody rabbit anti-PKMζ C-2 antisera (1:10,000) and chicken anti-EYFP (1:1,000) in BB. After washing 6 times for 10 mins each in PBS-T, the sections were then incubated with the secondary antibodies goat anti-chicken Alexa 488 and goat anti-rabbit-Alexa 647 (both 1:500 in BB; Jackson ImmunoResearch, West Grove, PA) for 2 h at room temperature. After extensive washing with PBS, the sections were mounted with 4′,6-diamidino-2-phenylindole (DAPI) Fluoromount-G (Southern Biotech, Birmingham, AL) or Vectashield (Vector Laboratories, Newark, CA).

Three sections from each mouse were observed using an upright Leica SP8 confocal microscope (Leica Biosystems, Buffalo Grove, IL) and analyzed using ImageJ (version 1.53a; 130). For each section, 8.5 µm-thick Z-stacks of the dorsal CA1, CA2, CA3, supra-DG, and infra-DG were created using the maximum intensity projection function in ImageJ. In each stack, two square regions of interest were centered on *strata pyramidale*, *radiatum*, *lacunosum moleculare* for the CA1, CA2, and CA3 subfields, and on the granule cell layer and molecular layer for both the supra- and infra-pyramidal layers of DG. The molecular layer of the DG was further subdivided into three equal compartments (inner, middle, and outer) along the dorsoventral axis. Measurements were made from each mouse in each region of interest (ROI). The raw integrated density (defined as the sum of the values for all pixels) of the Z-stack ROI expressing the fluorescent label was measured for the green (EYFP), red (PKMζ), and yellow (overlap) volume of pixels, and the average of each of the 6 (2 ROI x 3 slices) measurements was taken as representative for the region for each mouse. These measures account for the locations of the signal as well as their intensity. To rigorously compare the overlap between EYFP and PKMζ signals in the Trained and Untrained brain regions, we computed the Manders coefficients M1 and M2 using the JACOP plugin for ImageJ (131). M2 estimates how much PKMζ is in the memory-tagged cells by computing the proportion of the PKMζ signal that co-localizes with the EYFP signal, while M1 estimates the proportion of EYFP signal that co-localizes with the PKMζ signal. Based on prior work, the working hypothesis is that M2 will increase in persistently potentiated populations of synapses in somato-dendritic compartments that are maintained by a PKMζ-dependent mechanism (10, 16, 66).

### Preparation of micro-dissected hippocampal subfields (Experiment 2)

A day after the last training session, and 30 min after the retention test without shock, mice were anesthetized with 2% isoflurane and decapitated. Transverse 300-μm brain slices were prepared using a vibratome (model VT1000 S, Leica Biosystems, Buffalo Grove, IL) and incubated at 36°C for 30 min and then at room temperature for 90 min in oxygenated artificial cerebrospinal fluid (aCSF in mM: 125 NaCl, 2.5 KCl, 1 MgSO4, 2 CaCl2, 25 NaHCO3, 1.25 NaH2PO4, and 25 Glucose) as previously prepared to measure LTP and place avoidance memory-induced synaptic plasticity (10, 132). Slices were cut in half so that one hemisphere could be used for RNA-seq (see below) and one for *ex vivo* slice physiology (data not shown). The DG, CA3, and CA1 subfields were micro-dissected for later RNA extraction using a 0.25 mm punch tool and cold mat (Electron Microscopy Sciences, Hatfield, PA) under a dissecting scope (Zeiss, Oberkochen, Germany; Figure 2C).

### RNA-sequencing and quality control (Experiment 2)

RNA was isolated using the Maxwell 16 LEV RNA Isolation Kit (Promega, Madison, WI). Sequencing libraries were prepared by the Genomic Sequencing and Analysis Facility at The University of Texas at Austin and sequenced on the Illumina HiSeq platform following standard protocols. Reads were processed on the Stampede cluster at the Texas Advanced Computing Center (RRID#: SCR_021713). The quality of raw and filtered reads was checked using the program FASTQC (133) and visualized using MultiQC (134). We obtained 6.9 million ± 6.3 million reads per sample (Figure S1A). Next, we used Kallisto (135) to pseudo-align raw reads to a mouse reference transcriptome (Gencode version 7; 136), which yielded 2.6 million ± 2.1 million reads per sample (Figure S1B). Mapping efficiency was about 42% (Figure S1C).

### Statistical analyses

To enhance statistical power and reduce the impact of low-abundance noise, we first applied gene-level filtering prior to all statistical analyses. As the goals of differential expression and correlation analyses differ, we employed distinct pre-processing strategies optimized for each.

For differential expression analysis, we followed DESeq2 guidelines to filter low-count genes by retaining only those with a total count greater than 1 across all samples. Starting from 22,485 genes, we retained 16,992, 16,481, and 16,840 genes in the DG, CA3, and CA1 subfields, respectively. In the combined dataset, 17,911 genes remained. Variance-stabilizing transformation (VST) was applied prior to downstream analyses to improve statistical inference.

In contrast, correlation-based analyses prioritized the inclusion of genes showing meaningful variation. Gene expression values were normalized separately for each hippocampal subfield using sample-wise size factor adjustment based on gene-wise geometric means.

Genes were filtered to exclude those with minimal expression variation, retaining only those expressed above the minimum observed value in at least two samples. The final number of genes used for correlation analysis was 16,467 for DG, 15,778 for CA3, and 16,171 for CA1. Expression values were then log2-transformed to stabilize variance and avoid undefined values during correlation calculations.

All statistical analyses were performed using R version 4.0.0 (2020-04-24) (137), and Python 3.12.6. For behavioral feature analysis, we use the scikit-learn (v1.5.2; 138), scipy (v 1.14.1; 139) and Statsmodels (v0.14.4; 140) packages to conduct ANOVA and principal component analysis (PCA). One- and two-way ANOVAs were used to identify group differences in behavioral measures across one or multiple trials, respectively (141). PCA was conducted to reduce the dimensionality of the data (142, 143). In particular, to define PC1_mem_, we used behavioral features from the retention phase, as they provided the best description of memory performance. Transcript counts were imported into R and aggregated to yield gene counts using the gene identifier from the Gencode transcriptome. Differential expression analysis was performed using DESeq2 (144) with support from tidyverse packages (145, 146). Genes were considered significant at a false discovery rate (FDR) adjusted p-value threshold of < 0.1. Comparisons were made either among the four behavioral treatment groups (standard-trained, standard-yoked, conflict-trained, and conflict-yoked) or between the combined memory-trained and combined yoked-control groups.

Gene Ontology enrichment analysis was made using g:Profiler webserver/package (147) and MGI webserver (148) to identify biological terms associated with hypothesized and differentially expressed genes.

For analysis of non-linear associations between gene expression values, as well as to PC1_mem_ behavior, we defined Xicor (*ξ*) according to Chatterjee’s method (75). Since several methods exist for handling tied ranks, we tested multiple approaches and selected the ‘dense’ ranking method as default (Figure S5). Given our data limited sample size, we applied a correction to the observed Xicor (*ξ_observed_*) using a scaling model (Eq. 1) with sample size (*N*) and a scaling factor (*c*) derived from simulated linear data across varying sample sizes.

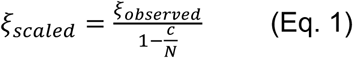

The newly generated Correlation Signal (Σ) from the norm of Spearman and Xicor correlation values follows Eq. 2.

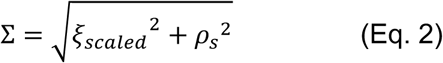

To calculate the p-value of Σ, we normalized both the observed Xicor (*ξ_observed_*)) and Spearman correlation, using their respective null variances:

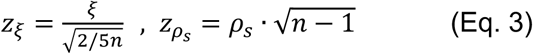

Assuming *z*_*ξ*_ and *z_ρs_* are independent standard normal variables, the combined statistic

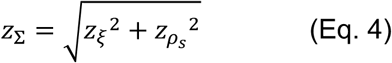

follows a Rayleigh distribution. The p-value was computed as the upper tail probability of the Rayleigh cumulative distribution function (CDF). Genes with p-values ≤ 0.05 were used for Louvain clustering.

Clustering and network analyses were performed through NetworkX (v3.4.2; 149) and python-louvain (v0.16; 150) and its visualization was generated in Cytoscape (v3.10.2; 151). WGCNA analysis was performed through WGCNA (ver1.69; 82) using the same genes used for the final clustering analysis. For visualization in R, we relied on the packages ggplot2 (145), cowplot (152), and png. We used Matplotlib (v3.9.2; 153) and Seaborn (v0.13.2; 154) packages in python. Final illustrations were created using Adobe Illustrator and Canvas X Draw.

## Data Availability

Raw sequence data and differential gene expression data are available in NCBI’s Gene Expression Omnibus Database (accession: GSE100225). All data, code, and results are publicly available on GitHub (https://github.com/sommerJY/APA_PKMZ).

## Acknowledgments

This work was initiated in the Neural Systems and Behavior course at the Marine Biological Laboratory. We thank the Promega Corporation for generously donating molecular supplies for RNA isolation. We thank Becca Young Brim, James Curley, Laura Sisk-Hackworth, Dennis Wiley, and Laura Colgin for comments on earlier versions of this manuscript. Supported by Chan Zuckerberg Initiative grant CP2-1-0000000614 (AAF/SM), and NIH grants R01MH132204 (AAF) and R01MH115304 (AAF/TCS). We are grateful to the Simons Center for Computational Physical Chemistry at NYU (Simons Foundation grant 839534, MT), Helmsley Foundation, a E.E. Just Endowed Research Fellowship, the Fries Trust Research Awards, Hartline MacNichol Research Awards, Great Generation Fund for Research, MGF Fuortes Memorial Research Awards, Frank R. Lillie Quasi-Endowment Fund, and L. & A. Colwin Summer Research Fellowship, and UT Austin Continuing Fellowship (RMH) that provided the support, without which these data could not be collected. This work was supported in part through the NYU IT High Performance Computing resources, services, and staff expertise.

## Author Contributions

Immunohistochemistry data analysis and analysis (AG-P), bioinformatic data analysis and presentation (JH, AP, SM), bioinformatic data curation and QC (RMH), data collection and analysis (H-YK, RMH, HAH, AAF), experimental design (HAH, JMA, AAF), figure preparation and data summary (JH, RMH), project management (SM, HAH, AAF), funding (SM, AAF), data interpretation (SM, HAH, AAF), and writing (AAF, SM, TCS, HAH).

