## Supplementary Figures and Tables for "Persistently increased expression of PKMzeta and unbiased gene expression profiles identify hippocampal molecular traces of a long-term active place avoidance memory and ‘shadow’ proteins"

### SUPPLEMENTARY INFORMATION

Table S1: Behavioral end-point measures of place avoidance.

Table S2: Discovery-driven analysis of GO terms that are positively correlated with $PC1_{mem}$.

Table S3: Analysis of GO terms hypothesized to play a role in learning and memory.

Figure S1. Quality control of RNA-sequencing data.

Figure S2. Experiment 2 number of datasets.

Figure S3. Astrocytic genes.

Figure S4. Robust detection of IEGs and DEGs that predict whether the sample was from the trained or control groups of mice using three non-linear methods.

Figure S5. Comparing the performance of Pearson, Spearman, and Xi correlations to analysis in detecting linear and non-linear co-expression relationships.

Figure S6. Impact of Correlation Methods and WGCNA Parameters on Module Detection

Figure S7. Gene Ontology analysis of the genes co-assigned with the three key genes by robust Louvain community detection.

Figure S8. PCA of the z-scored raw expression matrix fails to separate candidate genes or training conditions.

Figure S9. PCA visualization of raw Sigma(Σ) scores and TOM similarity.

Figure S10. Two-dimensional scatter plots of selected metric pairs.

Figure S11. Robustness of t-SNE and UMAP Dimensionality Reductions Across Parameter Settings.

Figure S12. Ablation study using different gene-gene correlation measurements.

Figure S13. Overall pipeline of the C-SCoRe.

| Variable | End-point Measure | Meaning |
|---|---|---|
| **TotalPath** | **Total Path** | **Total distance walked by the animal, in meters.** |
| **NumEntrances** | **Number of Entrances** | **Total number of times the animal entered the shock zone.** |
| **Time1stEntr** | **Time to first entrance** | **Amount of time it took the animal to enter the shock zone for the first time** |
| **Time2ndEntr** | **Time to second entrance** | **Amount of time it took the animal to enter the shock zone for the second time** |
| Time3rdEntr | Time to third entrance | Amount of time it took the animal to enter the shock zone for the third time |
| Time4thEntr | Time to fourth entrance | Amount of time it took the animal to enter the shock zone for the fourth time |
| Time5thEntr | Time to fifth entrance | Amount of time it took the animal to enter the shock zone for the fifth time |
| **Path1stEntr** | **Path to first entrance** | **Distance the animal walked before entering the shock zone for the first time** |
| **Path2ndEntr** | **Path to second entrance** | **Distance the animal walked before entering the shock zone for the second time** |
| Path3rdEntr | Path to third entrance | Distance the animal walked before entering the shock zone for the third time |
| Path4thEntr | Path to fourth entrance | Distance the animal walked before entering the shock zone for the fourth time |
| Path5thEntr | Path to fifth entrance | Distance the animal walked before entering the shock zone for the fifth time |
| **Speed1stEntr** | **Speed to first entrance** | **Speed at which the animal entered the shock zone for the first time** |
| **Speed2ndEntr** | **Speed to second entrance** | **Speed at which the animal entered the shock zone the second time** |
| **EntrPerDist** | **Entrances per distance** | **Number of entrances normalized by the total distance walked (#Ent/Tpath)** |
| **NumShock** | **Number of shocks** | **Total amount of shocks the animal received** |
| Timeto1stS | Time to first shock | Amount of time it took the animal to get shocked for the first time |
| Pathto1stS | Path to first shock | Distance the animal walked before getting shocked for the first time |
| **Speed** | **Speed** | **Average speed of the animal throughout the session** |
| **sdSpeed** | **SD of speed** | **Standard deviation of Speed** |
| **Linearity** | **Linearity** | **How straight or tortuous was the walking of the animal** |
| **MaxTimeAvoid** | **Maximum time avoided** | **Maximum amount of time that the animal stood away from the shock zone** |
| **MaxPathAvoid** | **Maximum path avoided** | **Maximum distance walked without entering the shock zone** |
| **TimeShockZone** | **Time in shock zone** | **Total time spent in the shock zone** |
| **pTimeShockZone** | **proportion in shock zone** | **Proportion of time spent in the quadrant where the shock zone is** |
| **pTimeCCW** | **Proportion in ccw** | **Proportion of time spent in the quadrant counterclockwise from the target quadrant** |
| **pTimeOPP** | **Proportion in opp** | **Proportion of time spent in the quadrant opposite from the target quadrant** |
| **pTimeCW** | **Proportion in cw** | **Proportion of time spent in the quadrant clockwise from the target quadrant** |
| **RayleigLength** | **Rayleigh length (A)** | **Length of the vector describing where the animal spent most of its time (the longer the vector, the stronger the preference for that spot. It ranges 0-1** |
| **RayleigAngle** | **Rayleigh angle (A)** | **Angle of the vector describing where the animal spent most of its time. It ranges 0-360.** |
| Polavg | Polar average (B) | Average degree where the animal spent most of time. It ranges 0-360. |
| PolSd | Polar SD | Standard deviation of polar average |
| PolMin | Polar minimum (C) | Proportion of time spent in the least preferred 10-degree bin |
| PolMinBin | Polar minimum bin (C) | The least preferred 10-degree bin |
| **Min50%RngLoBin** | **Minimum 50% low bin (D)** | **One of the boundaries that flank the least preferred arc (combination of least preferred bins)** |
| Min50HiBin | Minimum 50% high bin (E) | The other boundary that flanks the least preferred arc |
| PolMax | Polar maximum (F) | Proportion of time spent in the most preferred 10-degree bin |
| PolMaxBin | Polar maximum bin (F) | Most preferred bin |
| Max50LoBin | Maximum 50% low bin (G) | One of the boundaries that flank the most preferred arc (combination of most preferred bins) |
| Max50HiBin | Maximum 50% high bin(H) | The other boundary that flanks the most preferred arc |
| AnnMin | Annular minimum (I) | Proportion of time spent in the least preferred concentric circle |
| AnnMinBin | Annular minimum bin (J) | The least preferred concentric circle |
| AnnMax | Annular maximum (K) | Proportion of time spent in the most preferred concentric circle |
| AnnMaxBin | Annular maximum bin (L) | The most preferred concentric circle |
| Ann | Annular Average | Average annulus where the animal spent the time |
| AnnSD | Annular SD | Standard deviation of Annular average |
| **AnnularSkewnes** | **Annular Skewness** | **Skewness of the annular (histogram) distribution** |
| **AnnularKurtosis** | **Annular Kurtosis** | **Kurtosis of the annular (histogram) distribution** |
| **ShockPerEntrance** | **Shock per entrance** | **The number of shocks per entrance** |

**Table S1: Behavioral end-point measures of place avoidance.** The subset of measures used to quantify place avoidance are indicated by bold. The full set of raw behavioral data are available here: BehaviorRawData. The $PC1_{mem}$ and $PC2_{act}$ retention scores for each mouse are available here: BehaviorPCAdata.

| Domain | Name | Total | Present | Genes |
|---|---|---|---|---|
| MF | transcription regulator activity | 1789 | 50 | Npas4, Smad7, Fosl2, Nfil3, Btg2, Egr4, Srf, Ahr, Junb, Nr4a2, Atf3, Jun, Zfp281, Cited2, Nr4a3, Mtdh, Zbtb33, Egr1, Foxo1, Plagl1, Abhd2, Fosb, Rora, Dnajb1, Jund, Zfp869, Nr4a1, Zfp275, Klf2, Rfx2, Hspa1b, Maml2, Zfp655, Fos, Foxg1, Egr3, Ski, Jmjd1c, Erf, Bach1, Arid5b, Irf2bp2, Sox9, Jdp2, Pou3f3, Gmeb2, Myc, Med7, Kdm7a, Klf7 |
| MF | MAP kinase tyrosine/serine/threonine phosphatase activity | 13 | 6 | Dusp8, Dusp4, Dusp16, Dusp6, Dusp14, Dusp5 |
| MF | protein tyrosine/threonine phosphatase activity | 10 | 5 | Dusp8, Dusp4, Dusp16, Dusp6, Dusp5 |
| MF | molecular function regulator activity | 2135 | 39 | Rgs2, Ccnk, Smad7, Fbxw7, Sgk1, Bdnf, Scg2, Ranbp2, Eif5, Hmgcr, Irs2, Jun, Cited2, Rgs4, Flrt3, Ppp1r15a, Arpp19, Dnaja1, Apaf1, Rasl11a, Ccno, Dnajb4, Hsph1, Dnajb1, Jag1, Hspa1b, Calu, Ppp1r15b, Kitl, Ing2, Sh2d3c, Hsp90aa1, Rap2a, Il16, Cdkn1a, Arhgef3, Bag3, Tgfb3, Cpeb2 |
| MF | myosin phosphatase activity | 73 | 6 | Dusp8, Dusp4, Dusp16, Dusp6, Dusp14, Dusp5 |
| BP | learning or memory | 326 | 20 | Npas4, Ptgs2, Plk2, Arc, Nptx2, Adrb1, Pak6, Sgk1, Bdnf, Btg2, Kcnk10, Syt4, Srf, Nr4a2, Hmgcr, Jun, Egr1, Fos, Nedd9, Ntrk2 |
| BP | rhythmic process | 353 | 16 | Adrb1, Fbxw7, Nfil3, Bdnf, Ahr, Homer1, Per1, Jun, Fzd4, Adamts1, Egr1, Rora, Siah2, Ciart, Per2, Ntrk2 |
| BP | trans-synaptic signaling by neuropeptide | 3 | 3 | Bdnf, Syt4, Ntrk2 |
| BP | response to heat | 93 | 8 | Ptgs2, Arpp21, Dnaja4, Dnaja1, Dnajb1, Hspa1b, Hsp90aa1, Bag3 |
| BP | glucose metabolic process | 224 | 11 | Errfi1, Ranbp2, Irs1, Irs2, Atf3, Arpp19, Foxo1, Rora, Dyrk2, Serp1, Myc |
| BP | leukocyte activation | 1033 | 24 | Tnip2, Smad7, Fosl2, Peli1, Fzd5, Nfil3, Cxadr, Srf, Ahr, Irs2, Jun, Nr4a3, Egr1, Hsph1, Rora, Jund, Kitl, Egr3, Hsp90aa1, Cdkn1a, Irf2bp2, Arid1a, Myc, Nedd9 |
| BP | trans-synaptic signaling by BDNF, modulating synaptic transmission | 5 | 3 | Bdnf, Syt4, Ntrk2 |
| BP | cell migration involved in sprouting angiogenesis | 55 | 6 | Ptgs2, Plk2, Fbxw7, Srf, Nr4a1, Egr3 |
| BP | nuclear receptor-mediated glucocorticoid signaling pathway | 16 | 4 | Bdnf, Per1, Arid1a, Ntrk2 |
| BP | regulation of type B pancreatic cell proliferation | 17 | 4 | Errfi1, Irs2, Nr4a3, Nr4a1 |
| BP | response to amphetamine | 38 | 5 | Rgs2, Ranbp2, Nr4a2, Rgs4, Nr4a1 |
| BP | female pregnancy | 235 | 10 | Errfi1, Ptgs2, Rgs2, Junb, Cited2, Fosb, Fos, Arid1a, Slc2a1, Tgfb3 |
| BP | regulation of cell junction assembly | 236 | 10 | Fzd5, Bdnf, Flrt3, Lrrtm2, Amigo2, Fermt2, Rap2a, Slitrk5, Ntrk2, Vcl |
| BP | regulation of protein transport | 521 | 15 | Slc16a1, Ptgs2, Syt4, Hmgcr, Irs1, Irs2, Dnaja1, Foxo1, Plk3, Cyp51, Hsp90aa1, Serp1, Klf7, Bag3, Tgfb3 |
| BP | organic anion transport | 460 | 14 | Slc16a1, Ptgs2, Rgs2, Slc2a3, Bdnf, Syt4, Eprs, Irs2, Slc25a25, Rgs4, Slc6a6, Myc, Slc2a1, Ntrk2 |
| BP | vascular process in circulatory system | 238 | 10 | Ptgs2, Rgs2, Adrb1, Hmgcr, Klf2, C2cd4b, Gch1, Fermt2, Slc2a1, Dusp5 |
| CC | transcription factor AP-1 complex | 5 | 4 | Junb, Jun, Jund, Fos |
| CC | cell junction | 2266 | 42 | Npas4, Slc16a1, Smad7, Slc2a3, Pcdh8, Arc, Nptx2, Adrb1, Acan, Pak6, Fzd5, Bdnf, Kcnj2, Cxadr, Frmd6, Cul3, Cpeb4, Syt4, Eif5, Homer1, Flrt3, Mtdh, Fzd4, Kcna4, Dnajb1, Jag1, Mfap3l, Nr4a1, Lrrtm2, Fermt2, Egr3, Cldn12, Rap2a, Il16, Slitrk5, Slc6a6, Ctnnd1, Slc2a1, Gad1, Nedd9, Ntrk2, Vcl |
| CC | plasma membrane region | 1409 | 26 | Slc16a1, Ptgs2, Slc2a3, Pcdh8, Arc, Nptx2, Kcnj2, Cxadr, Frmd6, Irs1, Flrt3, Mtdh, Kcna4, Jag1, Hspa1b, Lrrtm2, Fermt2, Sh2d3c, Hsp90aa1, Rap2a, Arl13b, Slc6a6, Ctnnd1, Slc2a1, Nedd9, Ntrk2 |
| CC | cell-cell junction | 538 | 14 | Smad7, Fzd5, Kcnj2, Cxadr, Frmd6, Flrt3, Mtdh, Fzd4, Jag1, Fermt2, Cldn12, Ctnnd1, Slc2a1, Vcl |
| CC | neuronal dense core vesicle | 34 | 4 | Adrb1, Bdnf, Scg2, Syt4 |

**Table S2: Discovery-driven analysis of GO terms that are positively correlated with PC1.** Column 1 lists the three GO domains: Biological Process (BP), Cellular Compartment (CC), and Molecular Function (MF). Columns 2 lists the top 5 most significant functional categories that are represented in the list of 232 genes that are positively correlated with PC1, a measure of avoidance behavior.

| | Candidate Genes | DG DEGs | LTP Genes from Sanes & Lichtman 1999 |
|---|---|---|---|
| Response to stimulus (GO-0050896) | Fos, Fosl2, Npas4, Grin1, Gria1, Gria2, Pick1, Fmr1, Camk2a, Wwc1, Prkcb, Prkcz, Prkci | 2410002F23Rik, Abhd2, Adrb1, Ahr, Ankrd13c, Ankrd27, Apaf1, Arhgef3, Arid1a, Arid5b, Arl13b, Arpp21, Aste1, Atf3, B3gnt2, Bach1, Bag3, Bdnf, Btg1, Btg2, C2cd4b, Ccar2, Ccnk, Cdkn1a, Chrd, Cited2, Cpeb2, Cpeb4, Ctnnd1, Cul3, Cxadr, Cyp51, Dnaja1, Dnaja4, Dnajb1, Dusp16, Dusp4, Dusp5, Dusp6, Dusp8, Dvl2, Dyrk2, Egr1, Egr3, Errfi1, Fbxl2, Fbxw7, Fermt2, Flrt3, Fos, Fosb, Fosl2, Foxo1, Frmd6, Fzd4, Fzd5, Gadd45a, Gadd45g, Gch1, Gcnt2, Gpr19, Hmgcr, Hmgn2, Homer1, Hsp90aa1, Hspa1a, Hspa1b, Hsph1, Ifrd1, Il16, Ing2, Irs1, Irs2, Jag1, Jun, Junb, Jund, Kcnj2, Kctd12, Kdm6b, Kitl, Klf2, Klf7, Lbh, Lemd3, Lmna, Maml2, Mcl1, Mest, Mn1, Mtdh, Myc, Nedd9, Nfil3, Npas4, Nr4a1, Nr4a2, Nr4a3, Ntrk2, Nuak1, Odc1, Olfml2b, Pak6, Peg10, Peli1, Per1, Per2, Piga, Plk2, Plk3, Ppp1r15a, Ppp1r15b, Ptgs2, Pxn, Ranbp2, Rap2a, Rasd1, Rfx2, Rgmb, Rgs2, Rgs4, Rora, Scg2, Sde2, Sec61b, Serp1, Sgk1, Sh2d3c, Siah2, Ski, Slc16a1, Slc25a25, Slc2a1, Slitrk5, Smad7, Sox9, Srf, Tgfb3, Tiparp, Tnip2, Tra2b, Traip, Tsc22d2, Usp27x, Wdr45b, Zbtb33, Zfand5, Zfp207, Zfp668 | Ache, Adcy1, Adcy10, Adcy2, Adcy3, Adcy4, Adcy5, Adcy6, Adcy7, Adcy8, Adcy9, Adra1a, Adra1b, Adra1d, Adra2a, Adra2b, Adra2c, Adrb1, Adrb2, Adrb3, Bdnf, Cacna1a, Cacna1b, Cacna1c, Cacna1d, Cacna1e, Cacna1f, Cacna1s, Calb1, Calm1, Calm2, Calm3, Camk1, Camk4, Capn1, Capn10, Capn2, Capn3, Ccr7, Cd47, Cdh1, Cdh2, Chrm1, Chrm2, Chrm3, Chrm4, Chrm5, Chrna1, Chrna3, Chrna7, Chrnb1, Chrnb2, Chrnb3, Cnr1, Cnr2, Creb1, Dlg4, Drd1, Edn1, Efna5, Egf, Egr1, Egr2, Epha5, Erbb4, Fgf2, Fyn, Gabbr1, Gabra1, Gabra5, Gabrb1, Gabrb2, Gabrb3, Gap43, Gfap, Gria1, Gria2, Grin1, Grin2a, Grin2d, Grm1, Grm4, Grm5, Grm7, Gucy1a2, Gucy1b2, Gucy2c, Gucy2d, Gucy2e, Gucy2g, Homer1, Homer2, Homer3, Htr1a, Htr1b, Htr1f, Htr2a, Htr2b, Htr2c, Htr3a, Htr3b, Htr4, Htr5a, Htr5b, Htr6, Htr7, Il1b, Inhba, Itga1, Itga10, Itga11, Itga2, Itga2b, Itga3, Itga4, Itga5, Itga6, Itga7, Itga8, Itga9, Itgad, Itgae, Itgal, Itgam, Itgav, Itgax, Itgb1, Itgb1bp1, Itgb2, Itgb2l, Itgb3, Itgb4, Itgb5, Itgb6, Itgb7, Itgb8, Itgbl1, Itpka, Itpkb, Itpkc, L1cam, Mapk1, Mapk10, Mapk11, Mapk12, Mapk14, Mapk3, Mapk4, Mapk6, Mapk7, Mapk8, Mapk9, Mas1, Ncam1, Ngf, Nos1, Nos3, Nrg1, Nrg2, Nrg3, Nrgn, Ntrk2, Oprd1, Oprl1, Oprm1, Parp1, Phpt1, Pla2g10, Pla2g1b, Pla2g2a, Pla2g2d, Pla2g2e, Pla2g2f, Pla2g3, Pla2g4a, Pla2g4f, Pla2g5, Pla2g6, Pla2g7, Plat, Plcb1, Plcb2, Plcb3, Plcb4, Plcg1, Plcg2, Plg, Pnoc, Ppp3ca, Ppp3cb, Ppp3cc, Prkacb, Prkar1b, Prkcg, Prkcz, Prkg1, Ptn, Rab3a, Rarb, S100b, Serpine2, Sptbn1, Src, Stx1b, Syp, Th, Thy1, Tnc, Ube3a, Vamp2, Vamp3, Vamp4, Vamp8 |
| Translation (GO-0006412) | Fmr1 | Btg2, Cpeb2, Cpeb4, Eif5, Mrps31, Ppp1r15a, Ppp1r15b, Rgs2, Serp1 | Itga2, Mapk1, Mapk3 |
| Synapse organization (GO-0050808) | Npas4, Arc, Grin1, Gria1, Pick1, Numb | Amigo2, Arc, Bdnf, Chrd, Flrt3, Fzd5, Homer1, Lrrtm2, Nedd9, Npas4, Ntrk2, Pcdh8, Rap2a, Slitrk5 | Ache, Bdnf, Cacna1a, Cacna1s, Camk1, Cd47, Cdh1, Cdh2, Chrna1, Chrna7, Chrnb1, Chrnb2, Dlg4, Drd1, Efna5, Epha5, Erbb4, Fyn, Gabra1, Gabra2, Gabra3, Gabra5, Gabra6, Gabrb2, Gabrb3, Gap43, Gria1, Grin1, Grin2a, Grm5, Homer1, Htr1a, Htr4, Icam5, Itga3, Itgb1, Itgb3, Itpka, L1cam, Mapk14, Nrg1, Nrg2, Nrg3, Ntrk2, Ptn, Rab3a, Syn1, Tnc, Ube3a |
| Learning or memory (GO-0007611) | Fos, Npas4, Arc, Grin1, Gria1, Prkcz | Adrb1, Arc, Bdnf, Btg2, Chrd, Egr1, Fos, Hmgcr, Jun, Nedd9, Npas4, Nptx2, Nr4a2, Ntrk2, Pak6, Plk2, Ptgs2, Sgk1, Srf, Syt4 | Adcy1, Adcy3, Adcy8, Adra1b, Adrb1, Adrb2, Bdnf, Cacna1c, Cacna1e, Calb1, Camk4, Chrna7, Chrnb2, Cnr1, Creb1, Drd1, Egr1, Gabra5, Gabrb3, Gria1, Grin1, Grin2a, Grm4, Grm5, Grm7, Gucy2d, Htr2a, Htr2c, Htr6, Htr7, Il1b, Itga3, Itga5, Itga8, Itgb1, Ncam1, Ngf, Nrg1, Nrgn, Ntrk2, Oprl1, Pla2g6, Plcb1, Prkar1b, Prkcg, Prkcz, Ptn, S100b, Snap25, Th, Ube3a |

**Table S3: Analysis of GO terms hypothesized to play a role in learning and memory.** Column 1 contains four GO categories representing four hypotheses about which genes are important for memory formation and recall: 1) response to stimulus, 2) translation, 3) synapse organization, and 4) learning or memory. Column 2 contains the candidate genes described in the introduction. Column 3 contains genes that were differentially expressed in the dentate gyrus (DG) of trained and yoked control mice a day after memory training and thirty minutes after unreinforced expression of active place avoidance memory. Of those DEG, 149 are associated with hypothesis 1 that genes would be related to response to stimulus, which includes many immediate early genes that trigger responses in signaling cascades with membrane-bound receptors. Only 9 genes were associated with the hypothesis that translation machinery that executes the response are needed. Only fourteen genes are associated with "synaptic organization" that includes synaptic receptors and channels that express the response. Twenty genes associated with learning or memory at the behavioral level were differentially expressed. **Adrb1, Arc, Bdnf, Btg2, Chrd, Egr1, Fos, Hmgcr, Jun, Nedd9, Npas4, Nptx2, Nr4a2, Ntrk2, Pak6, Plk2, Ptgs2, Sgk1, Srf and Syt4**. All differentially expressed genes in the learning and memory list also overlap with the genes in the response to stimulus list. Column 4 contains the genes that were reviewed by Sanes and Lichtman, 1999 to play a role in long-term potentiation, a cellular model of memory. This list of 249 contains the reviewed genes and their isoforms. Only seven genes (shown in bold: **Adrb1, Bdnf, Egr1, Homer1, Ntrk2, Stmn4, and Vamp1**) from the Sanes and Lichtman list were found to be differentially expressed in the DG.

SUPPLEMENTARY FIGURES

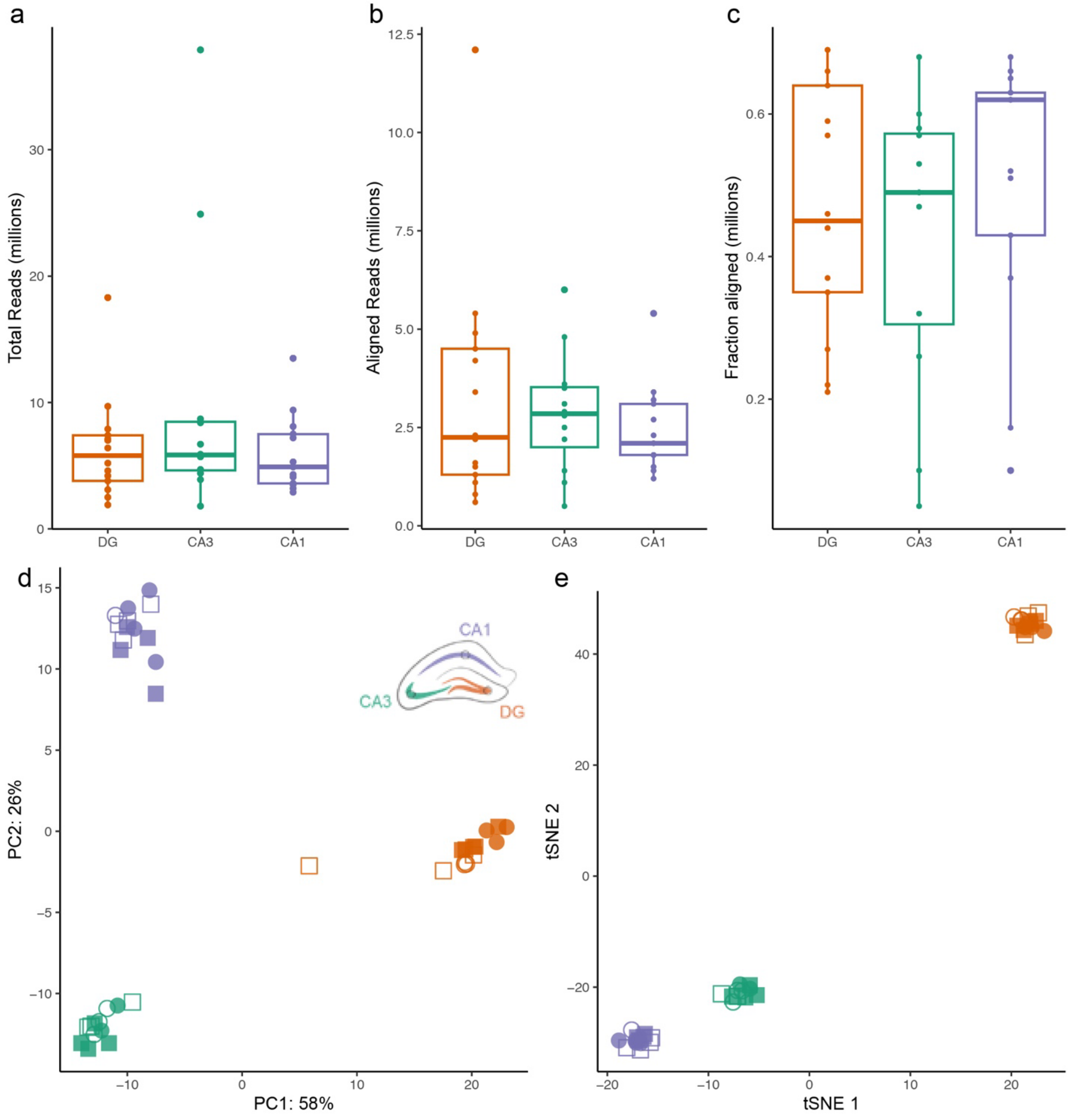


**Figure S1. Quality control of RNA-sequencing data.** a) On average, each sample yielded 6.9 million raw reads. b) Of those raw reads, on average 2.6 million mapped to the reference transcriptome. c) The alignment efficiency of Kallisto was over 40%. d) Principal component analysis (PCA) and e) t-Distributed Stochastic Neighbor Embedding (t-SNE) both confirm that all samples can be divided into three clusters based on subfield (DG, CA3, or CA1), as expected, with CA1 and CA3 being more similar to one another than to DG.

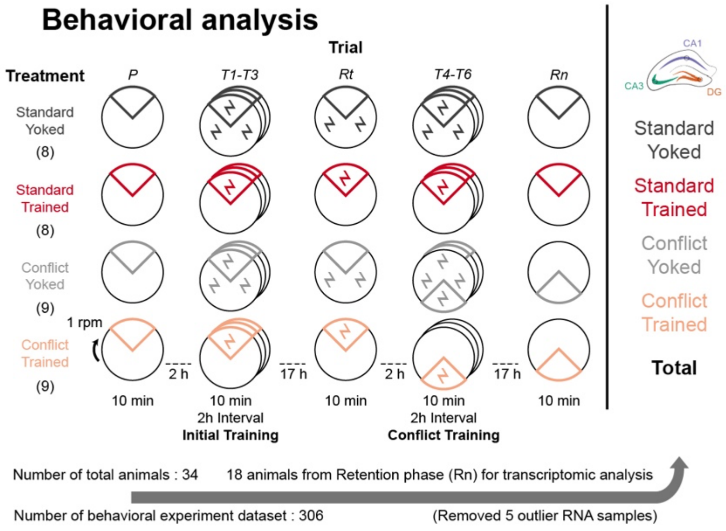


### RNA sequencing

| | Animal Count | DG | CA3 | CA1 | Sequenced Samples |
|---|---|---|---|---|---|
| Standard Yoked | 4 | 3 | 3 | 1 | 7 |
| Standard Trained | 4 | 3 | 2 | 4 | 9 |
| Conflict Yoked | 5 | 3 | 3 | 4 | 10 |
| Conflict Trained | 5 | 5 | 4 | 4 | 13 |
| **Total** | **18** | **14** | **12** | **13** | **39** |

Figure S2. Experiment 2 number of datasets.

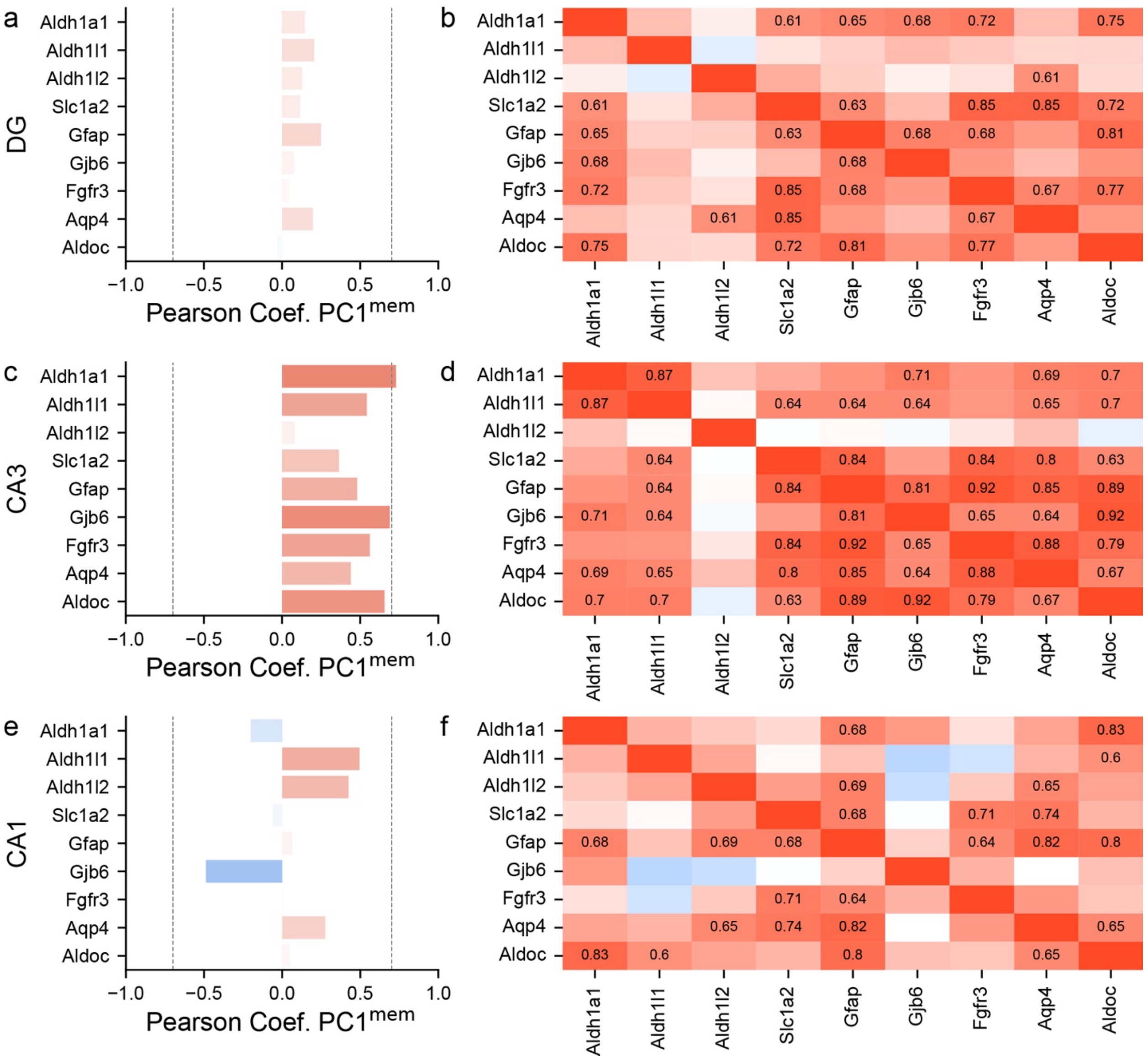


**Figure S3. Astrocytic genes**. Region-specific candidate gene analysis of astrocyte-related genes. None of the astrocyte-related genes were significantly correlated in DG (a) or CA1 (e), but *Aldh1l1, Aldoc, Fgfr*, and *Gfap* expression in the CA3 all show a correlation coefficient greater than 0.6 to behavioral PC1 (c). These candidate genes are highly, positively correlated with one another, especially in the CA3 (d) and to a lesser extent in the DG (b) and CA1(f).

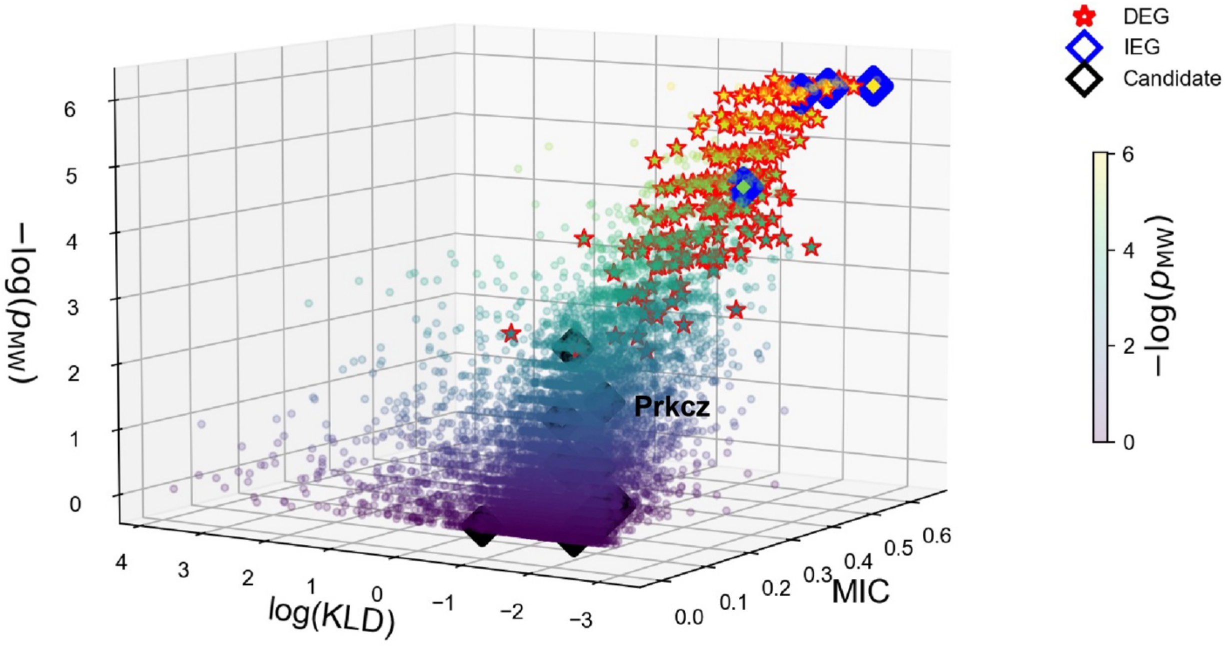


**Figure S4. Robust detection of IEGs and DEGs that predict whether the sample was from the trained or control groups of mice using three non-linear methods**. X: Kullback-Liebler Divergence (*KLD*) measured in bits. Y: Mutual Information (*MIC*) measured in bits. Z: Probability of the Mann-Whitney statistic ($p_{mw}$).

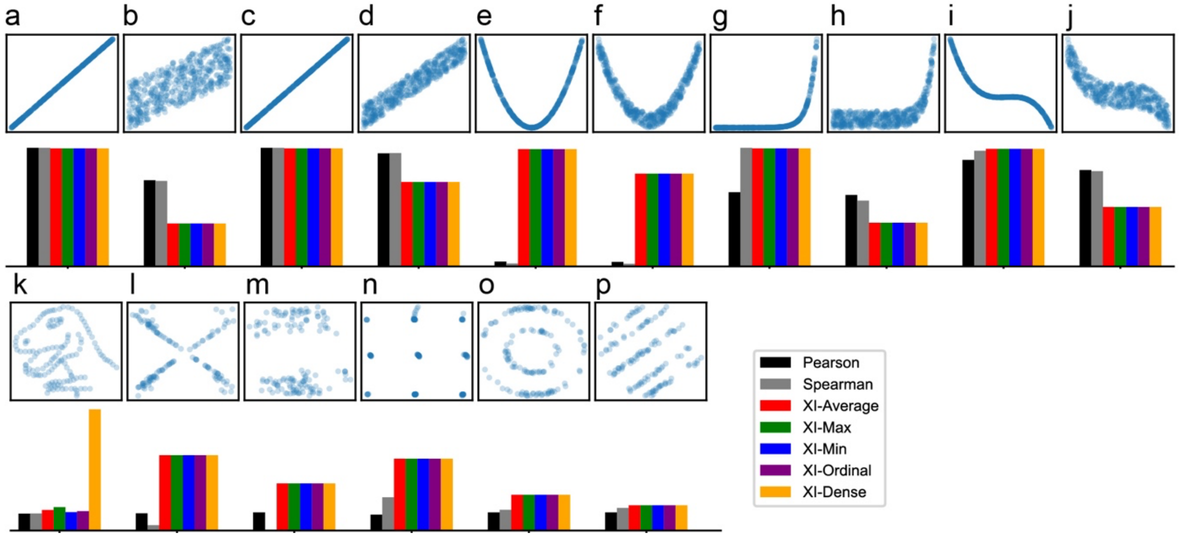


**Figure S5. Comparing the performance of Pearson, Spearman, and Xi correlations to analysis in detecting linear and non-linear co-expression relationships.**

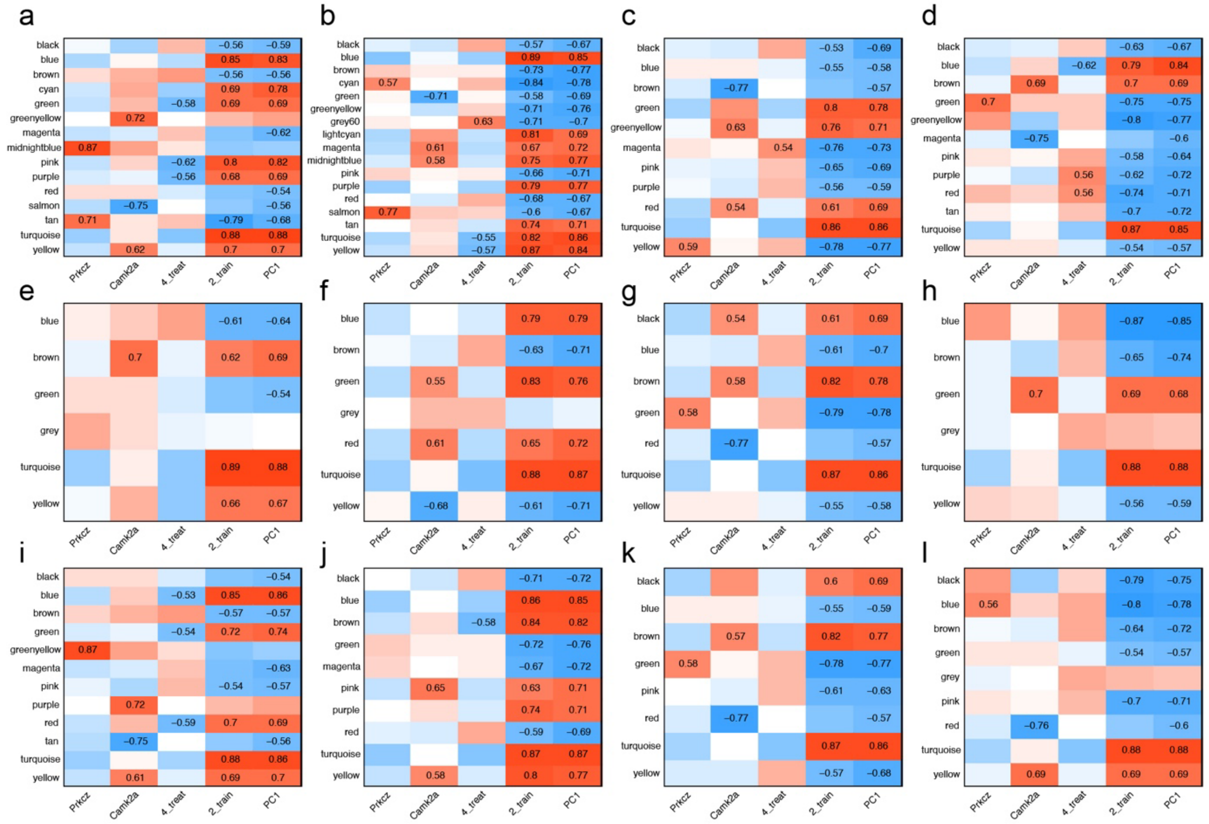


**Figure S6. Impact of Correlation Methods and WGCNA Parameters on Module Detection.** WGCNA was performed on 712 genes (excluding Prkcz and Camk2a to preserve them as external traits) using four gene–gene correlation matrices: Pearson (a, e, i), xi-correlation (b, f, j), sigma score (c, g, k), and max(Spearman, xi) (d, h, l). Module–trait relationships were evaluated against Prkcz expression, Camk2a expression, four treatment conditions, training status (yoked vs. trained), and $PC1_{mem}$. To assess parameter sensitivity, we varied deepSplit and minModuleSize across three settings: (a–d) deepSplit = 4, minModuleSize = 15, (e–h) deepSplit = 4, minModuleSize = 30, and (i–l) deepSplit = 2, minModuleSize = 15. These dramatic changes in module number and composition make it difficult to perform robust comparisons among the various gene–gene correlation methods.

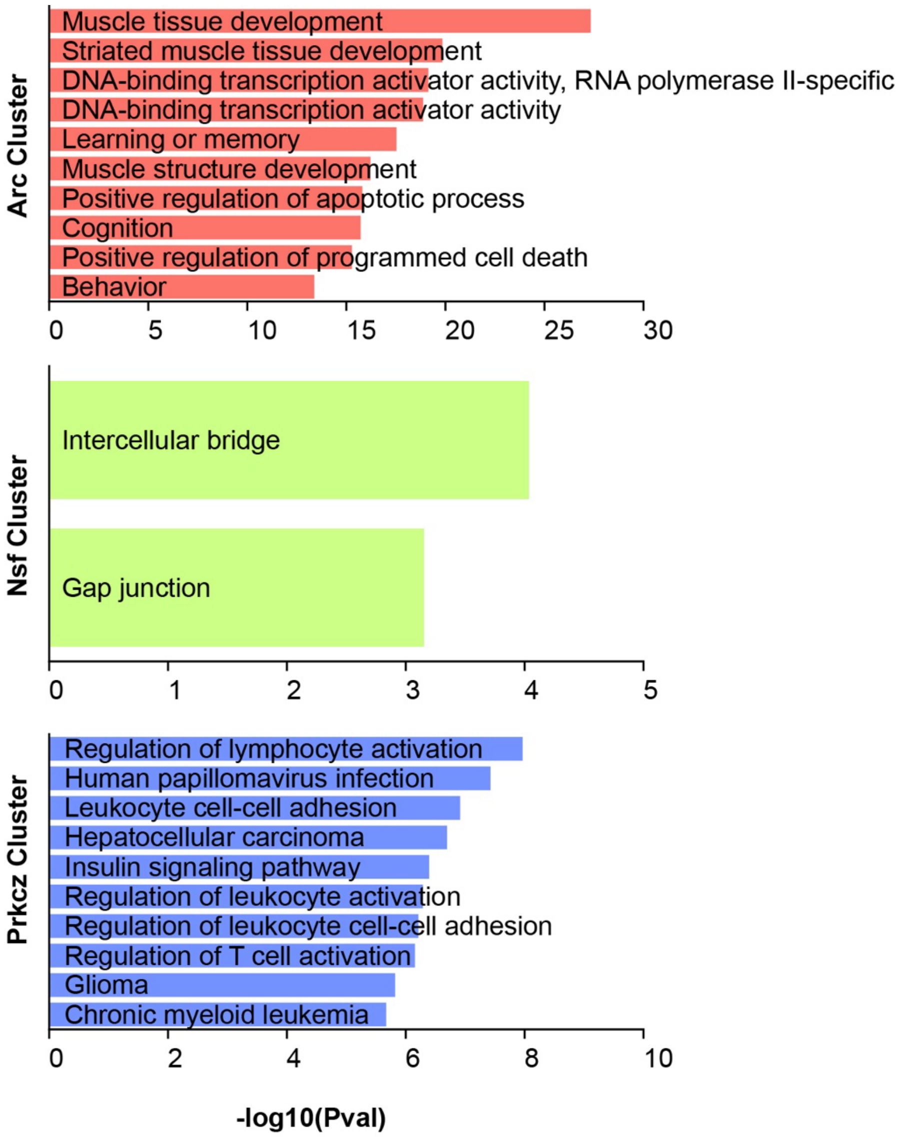


**Figure S7.** Gene Ontology analysis of the genes co-assigned with the three key genes by robust Louvain community detection**.**

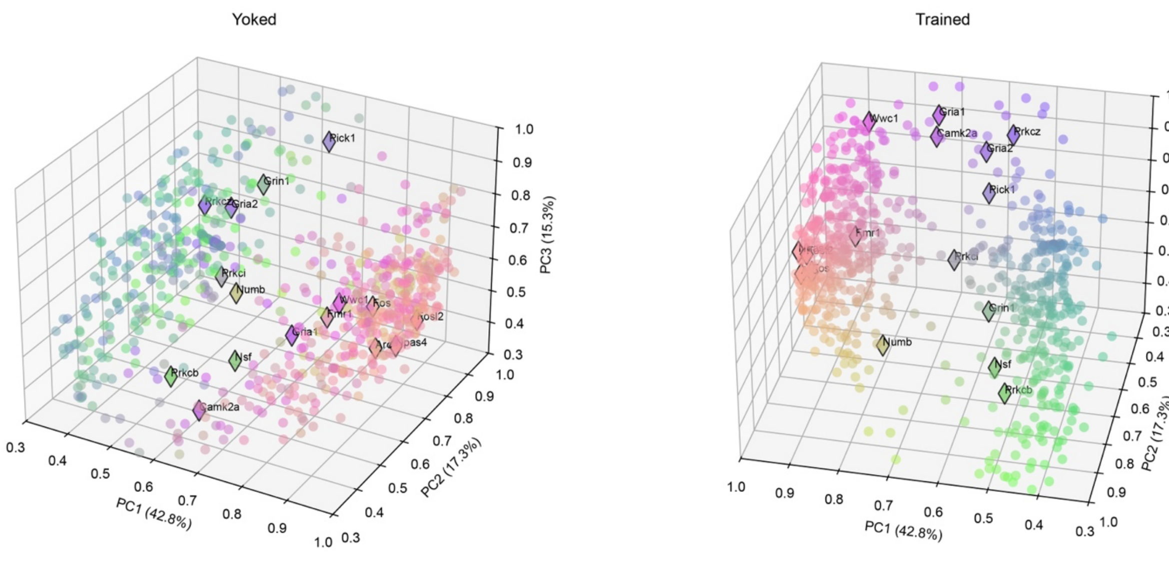


Figure S8. PCA of the z-scored raw expression matrix fails to separate candidate genes or training conditions.

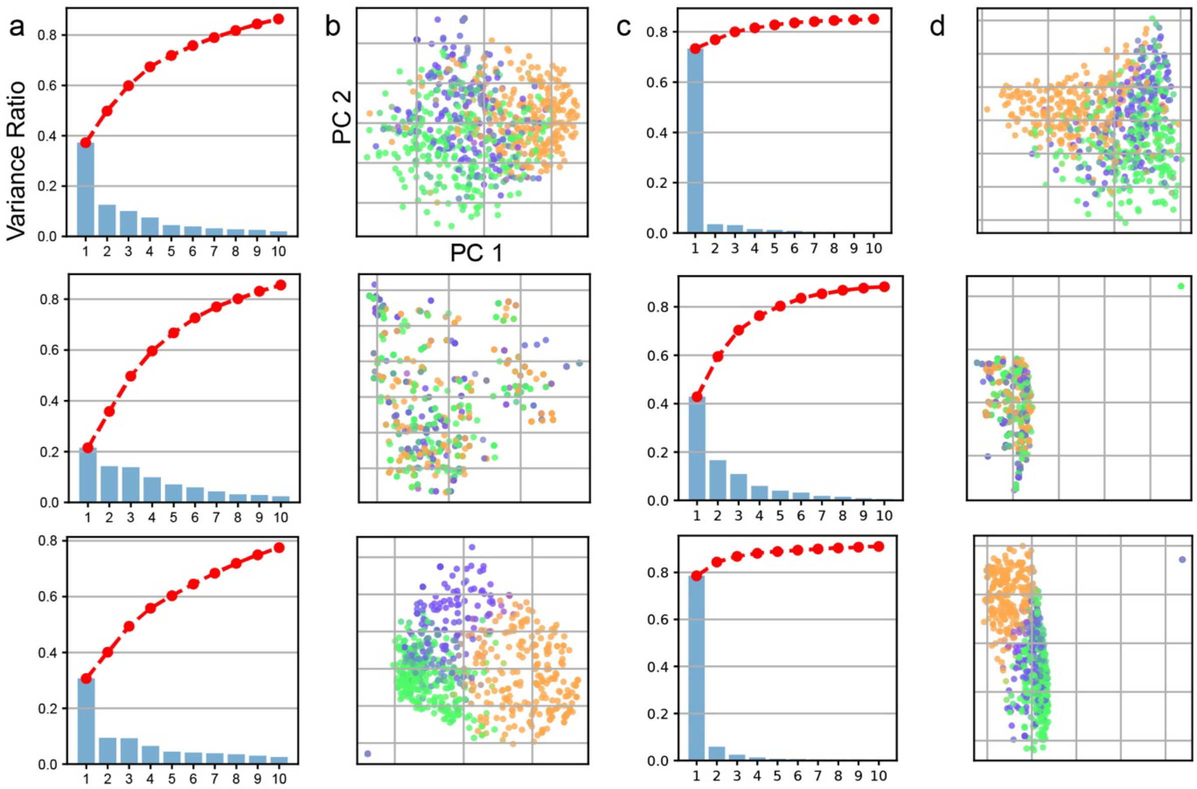


**Figure S9. PCA visualization of raw Sigma(Σ) scores and TOM similarity.** To compare with our gene–gene co-clustering score approach, we performed PCA using both the raw sigma correlation matrix (a, b) and the Topological Overlap Matrix (TOM) used in WGCNA (c, d). From top to bottom, each row represents all samples combined, yoked samples only, and trained samples only. We applied the same RGB color map as used for the main training clusters. These approaches tended to produce compact or ball-shaped clusters that made it less straightforward to distinguish condition-specific changes.

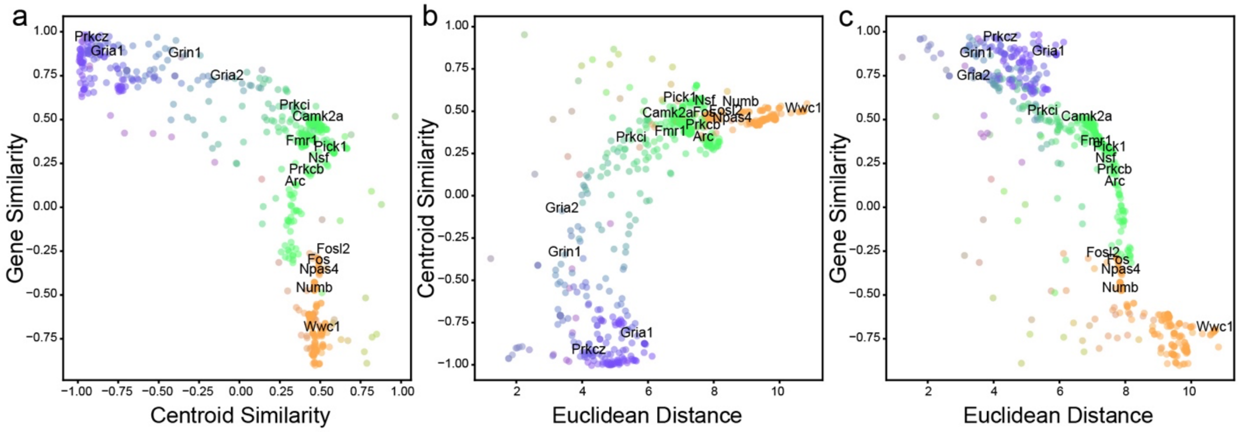


**Figure S10. Two-dimensional scatter plots of selected metric pairs.** These plots sufficiently illustrate the co-clustering trends observed in the data without requiring dimensionality reduction methods such as t-SNE or UMAP. The same RGB color map is used as for the main training clusters.

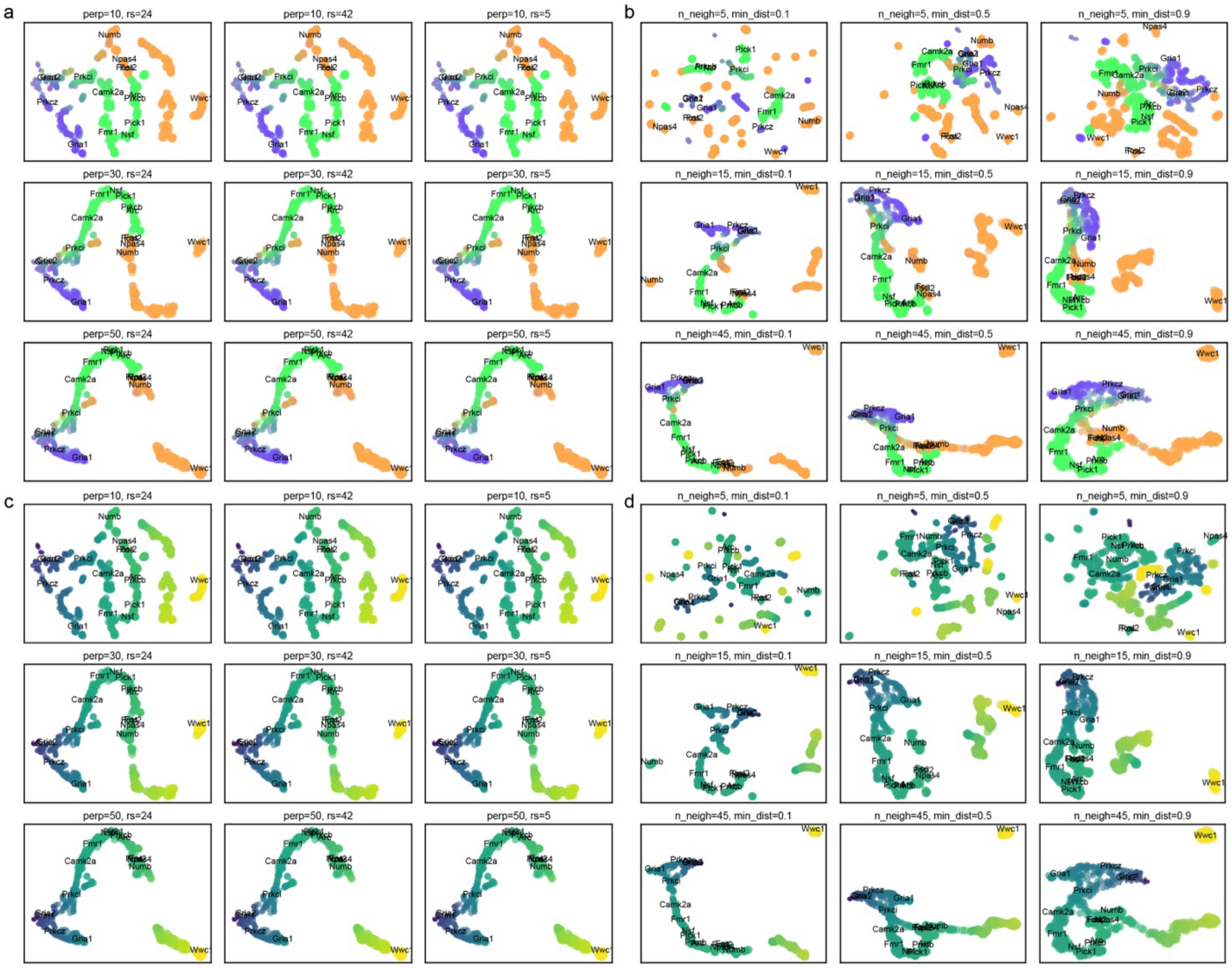


**Figure S11. Robustness of t-SNE and UMAP Dimensionality Reductions Across Parameter Settings.**
a) and c) t-SNE embeddings generated using varied perplexity (perp) and random seed (rs) settings. b) and d) UMAP embeddings generated using number of neighbors(n_neighbor) and minimum distances (min_dist). We used two coloring methods: 1) RGB color utilizing PC1, PC2 and PC3 (a, b) and 2) the color of the magnitude of Euclidean Distance changes from the yoked to the trained data set (c, d).

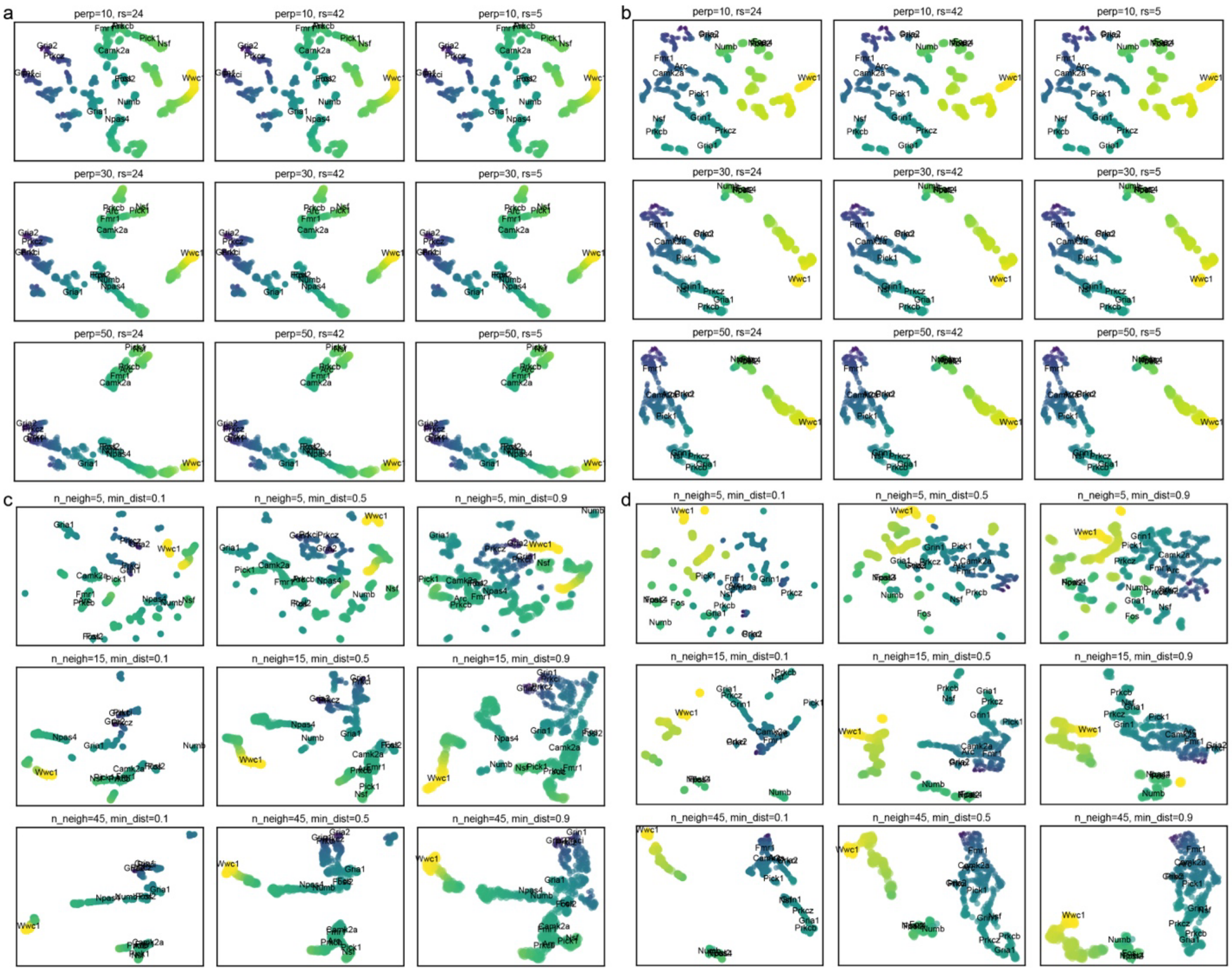


**Figure S12. Ablation study using different gene-gene correlation measurements.**

To test the robustness of the correlation method selection, we changed the gene-gene correlation measurements to the absolute value of Spearman correlation (abs(Spearman)) and Xi-Correlation (Xicor). The parameters varied are the same as the Figure S11. a) and c) are t-SNE and UMAP embedding results using abs(Spearman). b) and d) are results in the same order but using Xicor. The color represents the magnitude of Euclidean Distance from yoked to trained samples.

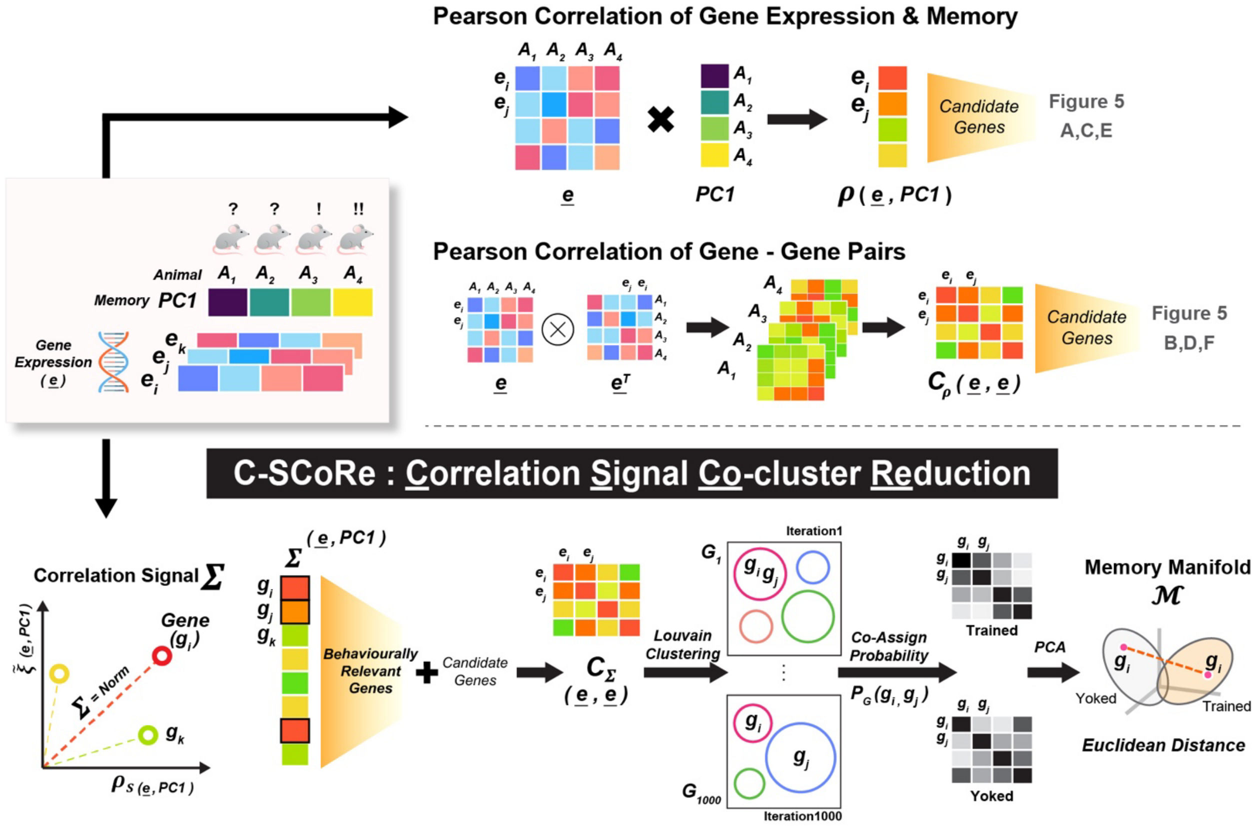


Figure S13. Overall pipeline of the C-SCoRe analysis.

The figure depicts the overall use of correlation measurements and pipeline in our manuscript. Above the dashed line shows the conventional usage of Pearson correlation on behavioral PC1-relevant genes and gene-gene pair interactions. The new method named C-SCoRe (Correlation Signal Co-cluster Reduction) is a series of procedures starting from calculation of the Correlation Signal and ending with the Co-Assign Probability (CAP) space reduction. $\mathrm{A}$ = Animal sample; $\underline{\mathrm{e}}$ = Overall gene expression matrix represented by gene-by-animal matrix; $\mathrm{e_i}$ = Gene expression value of gene $\mathrm{i}$; $\mathrm{PC1}$ = Memory measurement from behavioral PCA; $\mathrm{C_\rho}$ = Gene-Gene combination interaction score based on Pearson; $\rho_\mathrm{s}$ = Spearman correlation; $\tilde{\xi}$ = Normalized Xi-correlation score; $\mathrm{g_i}$ = Gene $\mathrm{i}$ indicator; $\Sigma$ = Correlation Signal score, which is the norm of $\rho_\mathrm{s}$ and $\tilde{\xi}$; $\mathrm{C}_\Sigma$ = Gene-Gene combination interaction score based on Correlation Signal ($\Sigma$); $\mathrm{G_1}$ = Group of subgraph results from Louvain clustering with seed number 1; $\mathrm{P_G(g_i, g_j)}$ = Probability of co-cluster assignment of gene $\mathrm{i}$ and gene $\mathrm{j}$ after the 1,000 iterations of Louvain clustering; $\mathcal{M}$ = Notation of memory manifold, which resulted from CAP dimension reduction; $\times$ = Inner product sign; $\otimes$ = Element-wise multiplication sign.
